# A geranylgeranyl reductase homolog required for cholesterol production in Myxococcota

**DOI:** 10.1101/2024.11.01.621555

**Authors:** Alysha K. Lee, Paula V. Welander

## Abstract

Myxococcota is a phylum of sterol producing bacteria. They exhibit a clade depth for sterol biosynthesis unparalleled in the bacterial domain and produce sterols of a biosynthetic complexity that rivals eukaryotes. Additionally, the sterol biosynthesis pathways found in this phylum have been proposed as a potential source for sterol biosynthesis in the last eukaryotic common ancestor, lending an evolutionary importance to our understanding of this pathway in Myxococcota. However, sterol production has only been characterized in a few species and outstanding questions about the evolutionary history of this pathway remain. Here, we identify two myxobacteria, *Minicystis rosea* and *Sandaracinus amylolyticus*, capable of cholesterol biosynthesis. These two myxobacteria possess a cholesterol biosynthesis pathway that differs in both the ordering and enzymes involved in biosynthesis compared to *Enhygromyxa salina*, a myxobacterium previously demonstrated to produce cholesterol, as well as the canonical pathways found in eukaryotes. We characterize an alternative bacterial reductase responsible for performing C-24 reduction, further delineating bacterial cholesterol production from eukaryotes. Finally, we examine the distribution and phylogenetic relationships of sterol biosynthesis proteins across both cultured and uncultured Myxococcota species, providing evidence for multiple acquisition events and instances of both horizontal and vertical transfer at the family level. Altogether, this work further demonstrates the capacity of myxobacteria to synthesize eukaryotic sterols but with an underlying diversity in the biochemical reactions which govern sterol synthesis, suggesting a complex evolutionary history and refining our understanding of how myxobacterial cholesterol production relates to their eukaryotic counterparts.

**Significance Statement:** Sterols are essential and ubiquitous lipids in eukaryotes, but their significance in bacteria is less understood. Sterol production in Myxococcota, a phylum of developmentally complex predatory bacteria, has provided insight into novel sterol biochemistry and prompted discussion regarding the evolution of this pathway within both the eukaryotic and bacterial domains. Here, we characterize cholesterol biosynthesis in two myxobacteria, providing evidence for distinct pathway organization and identifying a unique protein responsible for C-24 reduction. We couple these results to phylogenomic analysis of sterol biosynthesis within Myxococcota revealing a complicated evolutionary history marked by vertical and horizontal transfer, suggesting a mosaic acquisition of this pathway in Myxococcota and highlighting the complex role myxobacteria may have had in sterol transfer to eukaryotes.

## Introduction

While sterols are primarily thought of as eukaryotic lipids, a diverse and increasing set of bacteria capable of sterol biosynthesis have been discovered (1). Historically, sterol production within the bacterial domain was viewed as biosynthetically simple and attributed to horizontal transfer from eukaryotes (2, 3). However, Myxococcota, a phylum of social predatory bacteria (4–7), is unique among bacterial sterol producers for both the synthesis of eukaryotic-like sterols (8–10) and the depth of sterol production within the phylum. Further, previous studies characterizing sterol biosynthesis in this phylum have revealed a widespread capacity for novel biosynthesis enzymes. This includes the core pathways used to synthesize sterols, as exhibited by the unique variation on the bacterial C-4 demethylation enzymes used by *Enhygromyxa salina* to fully demethylate sterols (9) and extends to include downstream modifications, such as the novel steroid hydroxylation enzymes discovered in *Sorangium cellulosum* (11, 12). Additionally, myxobacteria produce steroids with modifications unaccounted for by our current knowledge of steroid biosynthesis enzymes (10, 13–15). Thus, Myxococcota displays a diversity of novel sterol biochemistry, providing a contrast to the canonical eukaryotic pathways. Yet, characterization of sterol production in Myxococcota has been limited to a few taxa suggesting that the potential for sterol synthesizers and novel sterol biochemistry within the phylum has not been fully explored.

Sterol biosynthesis in Myxococcota is also of interest for the insights it may provide into the evolutionary history of these lipids. Sterols are ancient lipids, and the last eukaryotic common ancestor (LECA) is thought to have possessed a pathway for complex sterol biosynthesis (3, 16–18). Additionally, sterols provide membranes with a flexibility that facilitates endocytosis (19–21) which may have been key in the endosymbiosis events underpinning eukaryogenesis. However, the origin of this pathway in LECA remains an open question. Phylogenomic analysis suggests a bacterial origin to sterol biosynthesis (22) and the phylogenetic relationship between sterol biosynthesis proteins in myxobacteria and eukaryotes indicates this origin may be in Myxococcota (23). These phylogenomic studies lend support to the syntrophy eukaryogenesis hypothesis, which leverages a tripartite syntrophic symbiosis between an ancient Asgard archaea, α-proteobacteria, and myxobacteria to explain the development of eukaryotic organelles and the high number of bacterial-like genes in the eukaryotic genome (24, 25). However, the discovery of fossilized sterols derived from simple biosynthesis pathways has suggested that eukaryotic sterol biosynthesis originated in stem group eukaryotes, and not in bacterial sterol producers. This has also raised questions about the timing of sterol acquisition in Myxococcota, the role of horizontal gene transfer in this acquisition, and their relevance to the ancient environments that played a backdrop for eukaryogenesis (17). But our understanding of Myxococcota ancestry broadly and, more specifically, the evolution of sterol synthesis within this phylum is limited as most studies are constrained to a few cultured species that belong primarily to the family *Myxococcaceae*. A deeper understanding of sterol synthesis, regulation, and function across the breadth of Myxococcota is necessary to begin to resolve these evolutionary questions.

In this study, we sought to further investigate complex sterol synthesis within Myxococcota. We identified two additional myxobacteria, *Sandaracinus amylolyticus* and *Minicystis rosea*, capable of de novo cholesterol biosynthesis. Genomic and lipid analyses revealed that the cholesterol biosynthesis pathway in these two myxobacteria is distinct from what we have previously observed in *E. salina*, as both *S. amylolyticus* and *M. rosea* are missing the canonical C-24 sterol reductase required for cholesterol production. Using a cell-free lysate system, we confirmed a bacterial C-24 reductase belonging to the geranylgeranyl reductase family is responsible for the saturation of the C-24 double bond on the sterol side-chain. This bacterial C-24 sterol reductase (Bsr) is found throughout the sterol producing families of Myxoccoccota as well as in the genomes of several cyanobacteria with the genetic capacity to produce sterols. However, Bsr is not present in the genomes of cholesterol-producing eukaryotes emphasizing that it is a cholesterol biosynthesis protein of bacterial origin. The unexpected discovery of a GGR homolog necessary for cholesterol production in Myxococcota prompted us to further investigate the distribution and phylogenomic context of other complex sterol biosynthesis proteins across Myxococcota, identifying likely instances of both vertical and horizontal transfer in the families which comprise this phylum. Through these analyses, we also show a capacity to produce sterols in uncultured myxobacteria and capture uncharacterized diversity in myxobacterial sterol synthesis. These results highlight Myxococcota as a unique bacterial source of the sterols typically associated with eukaryotes, providing additional examples of convergent evolution within bacterial sterol synthesis that underscore the biological importance and evolutionary complexity of this pathway across both the eukaryotic and bacterial domains.

## Results

### Myxobacteria from *Sandaracinaceae* and *Polyangiaceae* synthesize cholesterol

In our previous analysis of sterol biosynthesis genes in Myxococcota, we identified several myxobacteria with near complete cholesterol biosynthesis pathways (9). These myxobacteria belong to the families *Polyangiaceae* and *Sandaracinaceae* and are more distantly related to *Enhygromyxa salina*, which we have previously shown synthesizes cholesterol. To better clarify the biosynthetic potential of these bacteria, we first extracted free sterols from both *Sandaracinus amylolyticus* and *Minicystis rosea*. In extracts from both organisms, we identified cholesterol as well as C-24 saturated cholesterol intermediates (Fig. 1a; *SI Appendix*, Fig. S1). However, the relative proportions of these sterols vary between the two bacteria (*SI Appendix*, Fig. S2 and Table S1). The sterol profile of *S. amylolyticus* consists primarily of cholesterol and the intermediate lathosterol, with much lower concentrations of 7-dehydrocholesterol and several saturated sterols with spectra nearly identical to 7-dehydrocholesterol or lathosterol. Conversely, the *M. rosea* sterol profile consists primarily of the same saturated sterol intermediates we detected in *S. amylolyticus* but at higher relative concentrations. Additionally, neither of these bacteria produce any detectable C-24 unsaturated sterols, such as desmosterol or zymosterol, setting apart the sterol profile of these bacteria from *E. salina*, which is dominated by C-24 unsaturated sterols. The presence of only C-24 saturated intermediates suggests C-24 reduction occurs as an intermediary step in cholesterol biosynthesis (Fig. 1b) distinguishing cholesterol biosynthesis in these two families from *Nannocystaceae*, where C-24 reduction likely occurs as the terminal step in cholesterol biosynthesis (8, 9).

**Figure 1.**
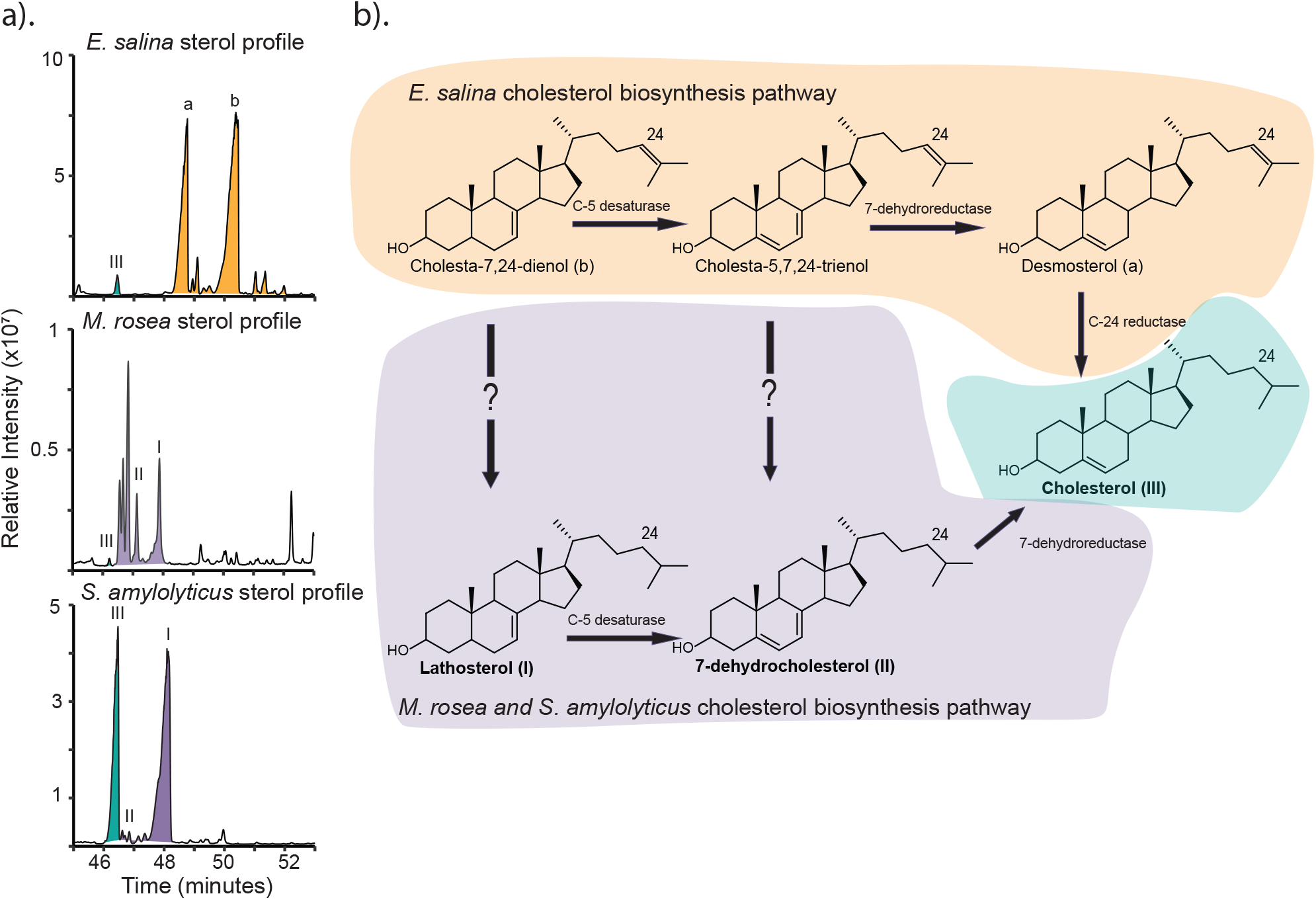
Sterol production in *Minicystis rosea* and *Sandaracinus amylolyticus*. (a) Total ion chromatograms of the sterols produced by *Sandaracinus amylolyticus* and *Minicystis rosea* with the sterols produced by *Enhygromyxa salina* for comparison. The sterol profile for both *S. amylolyticus* and *M. rosea* are characterized by C-24 saturated sterols, including cholesterol. None of the C-24 unsaturated sterols produced by *E. salina* are present in *S. amylolyticus* and *M. rosea*. Lipids were derivatized to trimethylsilyl ethers and the mass spectra for sterols identified are provided in Supplementary Figure 3.1. (b) Cholesterol biosynthesis pathway ordering differs between *E. salina* and *M. rosea* and *S. amylolyticus*, where *E. salina* performs C-24 reduction as the final step in biosynthesis with the canonical C-24 reductase and *M. rosea* and *S. amylolyticus* perform C-24 reduction at some intermediary step with an unknown biosynthesis enzyme.

Bioinformatic analyses revealed that both *S. amylolyticus* and *M. rosea* appear to have a cholesterol biosynthesis pathway that is largely homologous to the eukaryotic pathway, as we observed with *E. salina* (*SI Appendix*, Fig. S3). This includes the proteins involved in demethylation at C-14 and modification of the ring structure double bonds. However, *S. amylolyticus* and *M. rosea* lack the suite of eukaryotic proteins for C-4 demethylation and instead use the bacterial proteins, SdmAB, to fully demethylate at the C-4 position. This is also distinct from what we observed in *E. salina* which required a second reductase, SdmC, to fully demethylate at C-4. Cholesterol biosynthesis in *S. amylolyticus* and *M. rosea* further diverges from the eukaryotic pathway as neither bacterium has a homolog to the canonical C-24 reductase required to synthesize cholesterol or the C-24 saturated intermediates we observed, suggesting *S. amylolyticus* and *M. rosea* likely harbor a distinct enzyme for reducing the sterol side chain.

### Bacterial sterol C-24 reduction is carried out by a GGR family protein

To identify the enzyme responsible for C-24 reduction in these two myxobacteria, we first looked at the genomic context of known sterol biosynthesis genes in both organisms for potential candidates in the neighboring genes. Sterol biosynthesis genes in *S. amylolyticus* are largely organized into a single gene cluster (Fig. 2a). Among these genes, one coding for a geranylgeranyl reductase (GGR) family protein stood out as a candidate for C-24 sterol reduction. The GGR protein family (InterPro: IPR050407) includes a breadth of proteins from archaea, eukaryotes, and bacteria which all reduce isoprenoids of various chain lengths (26–29). Homologs to the *S. amylolyticus* GGR protein are found in species from all three sterol-producing families of Myxococcota and include *M. rosea* which also lacks the canonical C-24 reductase but produces C-24 saturated sterols *(SI Appendix*, Table S2). Additionally, we found homologs to this reductase in several cyanobacterial species, all of which have a genomic capacity to synthesize sterols (*SI Appendix*, Table S2). In the case of several of these bacteria, the GGR homologs also cluster with other sterol biosynthesis genes (*SI Appendix*, Fig. S4), further suggesting this protein may play a role in sterol biosynthesis. The distribution of this enzyme in sterol producing bacteria and its co-localization with known sterol biosynthesis genes prompted us to test it for C-24 reductase activity.

**Figure 2.**
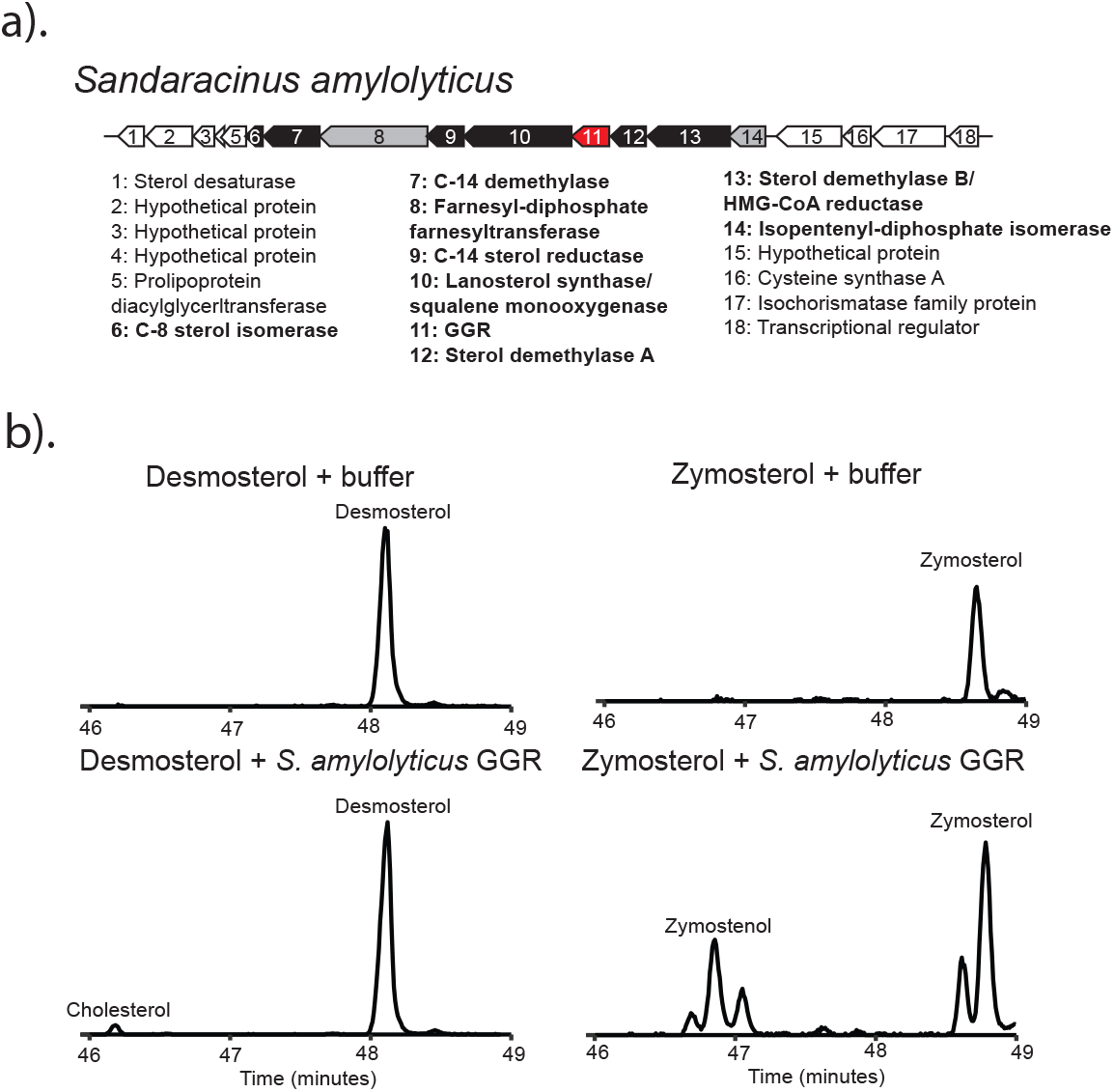
C-24 reduction in *S. amylolyticus* is carried out by a geranylgeranyl reductase (GGR) family protein. a) The sterol biosynthesis gene cluster in *S. amylolyticus*. Genes highlighted in black are involved in committed sterol biosynthesis steps. Genes highlighted in grey are involved in isoprenoid precursor biosynthesis. The gene highlighted red is a GGR homolog and putative bacterial C-24 sterol reductase (Bsr). b) Extracted ion chromatograms (*m/z*: 456, 458) of cell lysate experiments demonstrating saturation at C-24 by the *S. amylolyticus* Bsr protein. The substrates desmosterol or zymosterol were incubated with either lysis buffer or clarified *E. coli* cell lysate expressing Bsr. Sterol products were extracted through a modified Bligh Dyer.

We overexpressed the putative bacterial C-24 sterol reductase (Bsr) from *S. amylolyticus* in *Escherichia coli* to generate cell-free lysates. Because the *S. amylolyticus* sterol profile suggests C-24 reduction could be performed as an intermediary step in cholesterol biosynthesis, we tested both zymosterol and desmosterol as potential substrates for this protein. The *S. amylolyticus* GGR homolog converted zymosterol to zymostenol and desmosterol to cholesterol (Fig. 2b; *SI Appendix*, Fig. S5), demonstrating this enzyme is sufficient to perform C-24 sterol reduction. These data expand the known substrates of GGR family proteins to include sterols while also providing an alternative route for cholesterol production in bacteria.

The bacterial C-24 sterol reductase is found in all three sterol producing families of Myxococcota, suggesting this enzyme may be ancestral in the phylum. Because understanding the evolution of sterol biosynthesis within Myxococcota could provide important context for considering the evolution of sterol biosynthesis more broadly, we were interested in further probing the phylogenetic relationships between these GGR homologs. We constructed a maximum likelihood tree of GGR family proteins (Fig. 3a; *SI Appendix*, Fig. S6) including the Bsr homologs. The Bsr sequences all cluster together with the menaquinone reductases found in Mycobacteriales. This clade is distinct from the archaeal homologs involved in membrane biosynthesis, the bacterial homologs responsible for resistance to hydroxylamine, and the photosynthetic homologs involved in chlorophyll biosynthesis. We then compared the branching pattern of the bacterial C-24 sterol reductase homologs to a phylogenetic reconstruction of 16S sequences from cultured sequenced Myxococcota isolates as a proxy for taxonomy. Despite its distribution across the families of Myxococcota, the bacterial C-24 sterol reductase from this phylum does not mirror 16S based taxonomy (Fig. 3b). Bsr sequences belonging to both *Polyangiaceae* and *Nannocystaceae* branch separately and the sequences *Polyangiaceae* and *Sandaracinaceae* and sister to *Myxococcaceae* instead of *Nannocystaceae*. While this bacterial C-24 sterol reductase presents a unique example of convergent evolution within myxobacterial sterol biosynthesis, this observed branching pattern is inconsistent with vertical inheritance, suggesting horizontal transfer may have played a role in the proliferation of this enzyme in the phylum and leaving open the question of how ancestral sterol biosynthesis is in Myxococcota.

**Figure 3.**
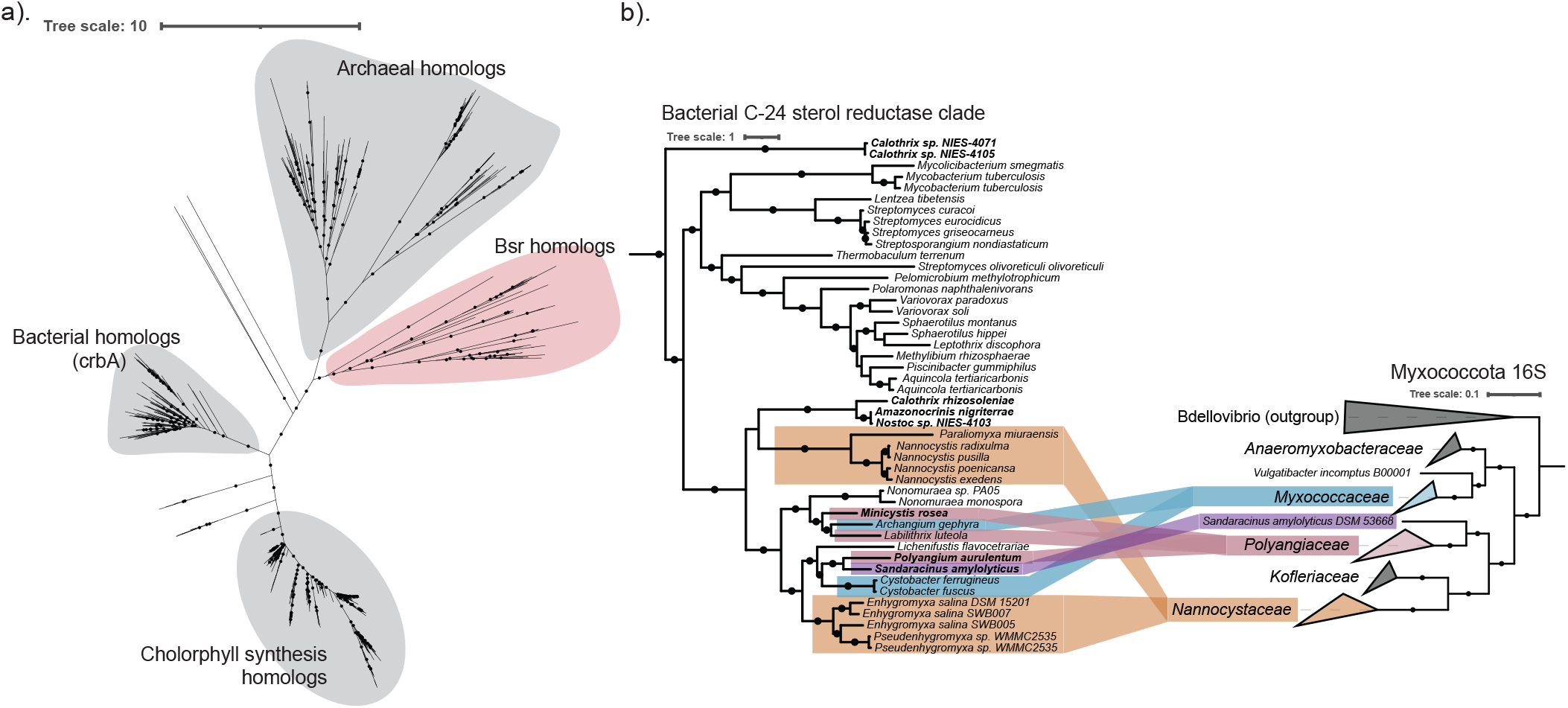
Bsr is a GGR homolog restricted to bacteria. a) An unrooted maximum likelihood tree of GGR homologs was generated using IQTree with the model of best fit and 5000 ultrafast bootstrap replicates. Branches with bootstrap support values >90 are denoted with a black circle. The clade containing the bacterial C-24 sterol reductase is highlighted in red. b) Expansion of the Bsr clade from the GGR tree shown in (a). Myxobacterial sequences are colored by their respective families as follows: *Nannocystaceae* (orange), *Sandaracinaceae* (purple), *Polyangiaceae* (pink), and *Myxococcaceae* (blue). Bolded species represent those with Bsr homologs co-localized with known sterol biosynthesis genes.

### Phylogenetic analysis of sterol biosynthesis across Myxococcota

Our findings demonstrate that proteins involved in complex sterol synthesis are found across Myxococcota but with an inherent complexity that does not always reflect vertical transfer of this pathway through the phylum. To further explore the evolution of sterol biosynthesis within this phylum, we expanded our phylogenetic analysis beyond Bsr by comparing the tree topology of other sterol biosynthesis proteins to the tree topology of Myxococcota 16S sequences. In doing so, we identify instances where these proteins have been vertically transferred, reflected by a congruence with myxobacterial 16S based taxonomy, and parts of the pathway that were inherited together, indicated by similar branching patterns across the biosynthesis protein trees.

*Myxococcaceae* is the most well studied family of Myxococcota, making up a majority of the cultured genomes we considered here. Many of these isolates harbor an oxidosqualene cyclase (Osc) homolog which is responsible for the initial cyclization of oxidosqualene to the first products of sterol biosynthesis – lanosterol or cycloartenol (30). These Osc homologs co-localize with squalene monooxygenase (Smo) in the genome and all are predicted to synthesize cycloartenol, in line with the sterol analysis of both *Stigmatella* and *Cystobacter* species (1, 8). The Osc from *Myxococcaceae* cluster together in a monophyletic clade that recapitulates 16S phylogeny (Figure 3). This branching pattern suggests Osc has been vertically transferred through the family and is ancestral to *Myxococcaceae*. This Osc homolog is not found in either of the families more closely related to *Myxococcaceae* and instead found only in one other myxobacterium, belonging to the family *Polyangiaceae*. With the exception of the bacterial C-24 sterol reductase homologs found in *Cystobacter* species, the proteins required for downstream modifications of the core sterol structure are absent from this family suggesting these myxobacteria do not further modify cycloartenol.

*Nannocystaceae* is a much smaller and understudied family of Myxococcota. All cultured, sequenced members of *Nannocystaceae* have an Osc homolog but unlike *Myxococcaceae* this Osc homolog synthesizes lanosterol (Fig. 4; *SI Appendix*, Fig. S7). Additionally, Osc does not co-localize with squalene monooxygenase in the genomes of *Nannocystaceae*. Osc sequences from this family form a monophyletic clade congruent with 16S phylogeny, suggesting Osc was vertically inherited. This clade branches separate from *Myxococcaceae* and this, alongside the differences in Osc type within these two families, suggests at least two distinct acquisitions of Osc in Myxococcota. *Nannocystaceae* also harbors myxobacteria that produce complex sterols (1, 8, 9, 31), and all members of this family have the required proteins for both C-14 and C-4 demethylation, indicating this capacity is widespread. The proteins for C-14 and C-4 demethylation as well as the canonical C-24 reductase cluster monophyletically in their respective phylogenetic trees, again recapitulating 16S phylogeny (*SI Appendix*, Fig. S8-S11). The branching patterns we observe for the sterol biosynthesis proteins in this family are conserved across these trees, suggesting sterol demethylation proteins were acquired with Osc and are also vertically inherited within the family. *E. salina* is the only member of *Nannocystaceae* to have all the proteins for ring structure double bond modifications required for cholesterol production (*SI Appendix*, Fig. S12). The remaining desaturation proteins are found in some of the isolates of *Nannocystaceae*, with those more closely related to *E. salina* exhibiting a more complete cholesterol biosynthesis pathway. This limited distribution could be the product of either gene loss or a later acquisition as the family diverged.

**Figure 4.**
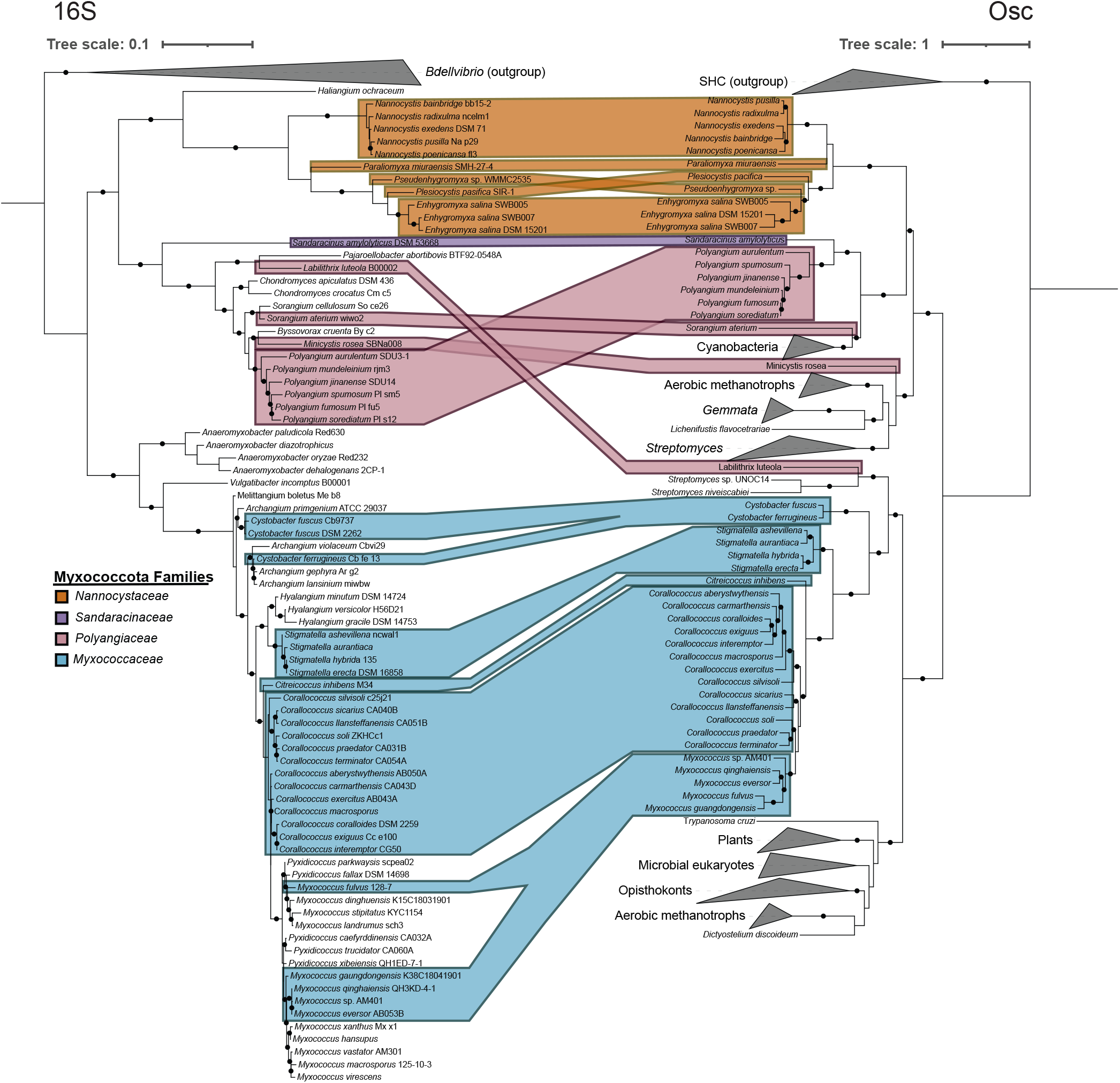
Phylogenetic analysis of myxobacterial oxidosqualene cyclase sequences (Osc) in relation to taxonomic diversity. A maximum likelihood trees of 16S sequences from Myxococcota species with *Bdellvibrio* sequences as an outgroup and Osc sequences with squalene hopane cyclase (SHC) sequences as an outgroup were generated using IQTree with the model of best fit and 5000 ultrafast bootstrap replicates. Branches with bootstrap support values >90 are denoted with a black circle. Myxobacterial genera are mapped from the 16S to Osc tree by boxes and these boxes are colored by their respective families as follows: *Nannocystaceae* (orange), *Sandaracinaceae* (purple), *Polyangiaceae* (pink), and *Myxococcaceae* (blue).

*Polyangiaceae* and *Sandaracinaceae* are also underrepresented and understudied families of Myxococcota. Osc is more sparsely distributed in these families (Fig. 4; *SI Appendix*, Fig. S7). With the exception of *Labilithrix luteola*, these Osc sequences are all lanosterol synthases, however, these sequences display a more complicated branching pattern than those found in the other families of Myxococcota. Osc sequences from *Polyangium* species cluster monophyletically and sister to the Osc sequence from the single cultured representative of *Sandaracinaceae*, recapitulating 16S phylogeny and perhaps suggesting Osc has been vertically transferred in these families as well. Interestingly, the remaining Osc sequences do not branch with *Polyangiaceae*. The *Sorangium aterium* Osc sequence clusters with the Cyanobacterial Osc sequences, and the *Minicystis rosea* Osc sequence clusters with other bacterial sequences. We posit that the placement of these branches, along with the cycloartenol synthase in *L. luteola*, represents multiple horizontal transfer events of Osc either into or out of *Polyangiaceae*.

Additionally, while we have identified several myxobacteria within these families capable of complex sterol production, the genes required for this process are even more sparsely distributed than Osc. The three isolates with a C-14 demethylase (Cyp51) do not branch together. The sequence from *Polyangium aurulentum* is sister to the sequences from *Nannocystaceae* and this may be in line with vertical transfer from an ancestor of *Polyangiaceae* and *Nannocystaceae* or transfer from an ancestor of *Nannocystaceae* to *P. aurulentum*. The sequences from *M. rosea* and *S. amylolyticus* branch separately from the other myxobacterial sequences and are basal to Cyp51 sequences from methanotrophs and sterol degrading pathogens (*SI Appendix*, Fig. S8 and S9). The placement of these branches again speaks to a possibility for horizontal transfer either into or out of these families. Additionally, the *S. amylolyticus* and *P. aurulentum* Cyp51 homologs do not branch together unlike their Osc sequences, raising the possibility that at least some of the sterol biosynthesis enzymes in *S. amylolyticus* exhibit a distinct evolutionary history. We also analyzed the phylogeny of bacterial C-4 demethylation proteins in these families however, these trees were not as robust as Cyp51, with sequences from *Polyangiaceae* and *Sandaracinaceae* branching with low support and clustering with different sequences depending on the model used (*SI Appendix*, Fig. S10).

Finally, the proteins involved in desaturation modifications are absent from most sterol producing members of these families, except for *P. aurulentum, M. rosea* and *S. amylolyticus* which have all the genes required for cholesterol production (*SI Appendix*, Fig. S12). This would suggest that most isolates from *Polyangiaceae* with a capacity for sterol production make biosynthetically simple sterols. This is particularly noteworthy in the *Polyangium* species, where *P. aurulentum*, a species basal to all other *Polyangium* isolates, has an apparent pathway for cholesterol production and the remaining *Polyangium* isolates have only an Osc homolog. This suggests that either *P. aurulentum* acquired downstream genes for cholesterol production after diverging from the other *Polyangium* species or that these *Polyangium* species lost the capacity for cholesterol production but maintained the ability to produce sterols. A similar pattern of either pathway loss across most members of *Polyangiaceae* or multiple acquisitions of cholesterol biosynthesis likely played out across this family. Altogether, these analyses suggest complex sterol biosynthesis is a trait restricted to the deep branching members of Myxococcota. However, the evolution of complex sterol biosynthesis is not consistent across the families of this phylum, with evidence of vertical transfer of key proteins in these pathways in some species but not all. As we continue to culture and study the species from these underrepresented families, the evolutionary relationships underpinning the presence of this pathway in Myxococcota may be better resolved.

### Myxococcota sterol biosynthesis proteins from metagenomes mirror the phylogenetic patterns observed in cultured isolates

Most of the diversity in Myxococcota remains uncultured (32, 33). To account for how these uncultured representatives might reshape our understanding of sterol biosynthesis in the phylum, we further extended our analysis of sterol biosynthesis proteins to include metagenome assembled genomes (MAGs) assigned to Myxococcota. We identified 121 unique Osc sequences from Myxococcota MAGs (Fig. 5). These MAGs span a broad range of ecosystems and include the terrestrial and marine environments where isolates have been previously cultured, as well as environments where myxobacteria have yet to be isolated, such as hydrothermal and host-associated systems. Many of these sequences fall into clades with cultured Myxococcota sequences but often on long branches, illustrating further diversity in sterol producing myxobacteria not currently captured in the cultured isolates. When assigned a more specific taxonomic ranking, sequences belonging to *Nannocystaceae* and *Myxococcaceae* cluster with cultured representatives from the same family, reflecting what we observed with cultured Osc sequences and further suggesting sterol biosynthesis may be ancestral in these families. Sequences from uncultured *Polyangiaceae* and *Sandaracinaceae* cluster throughout the tree. Some of these sequences branch with the cultured isolates while others branch with other bacterial or eukaryotic sequences. This broad branching pattern may again be evidence of horizontal transfer of sterol biosynthesis proteins to specific lineages in each family. Furthermore, additional Osc sequences did not affect the topology of the cultured *Polyangiaceae* sequences that cluster with the other bacterial Osc sequences, supporting their placement in the tree and providing evidence for horizontal transfer between these lineages and members of *Polyangiaceae*.

**Figure 5.**
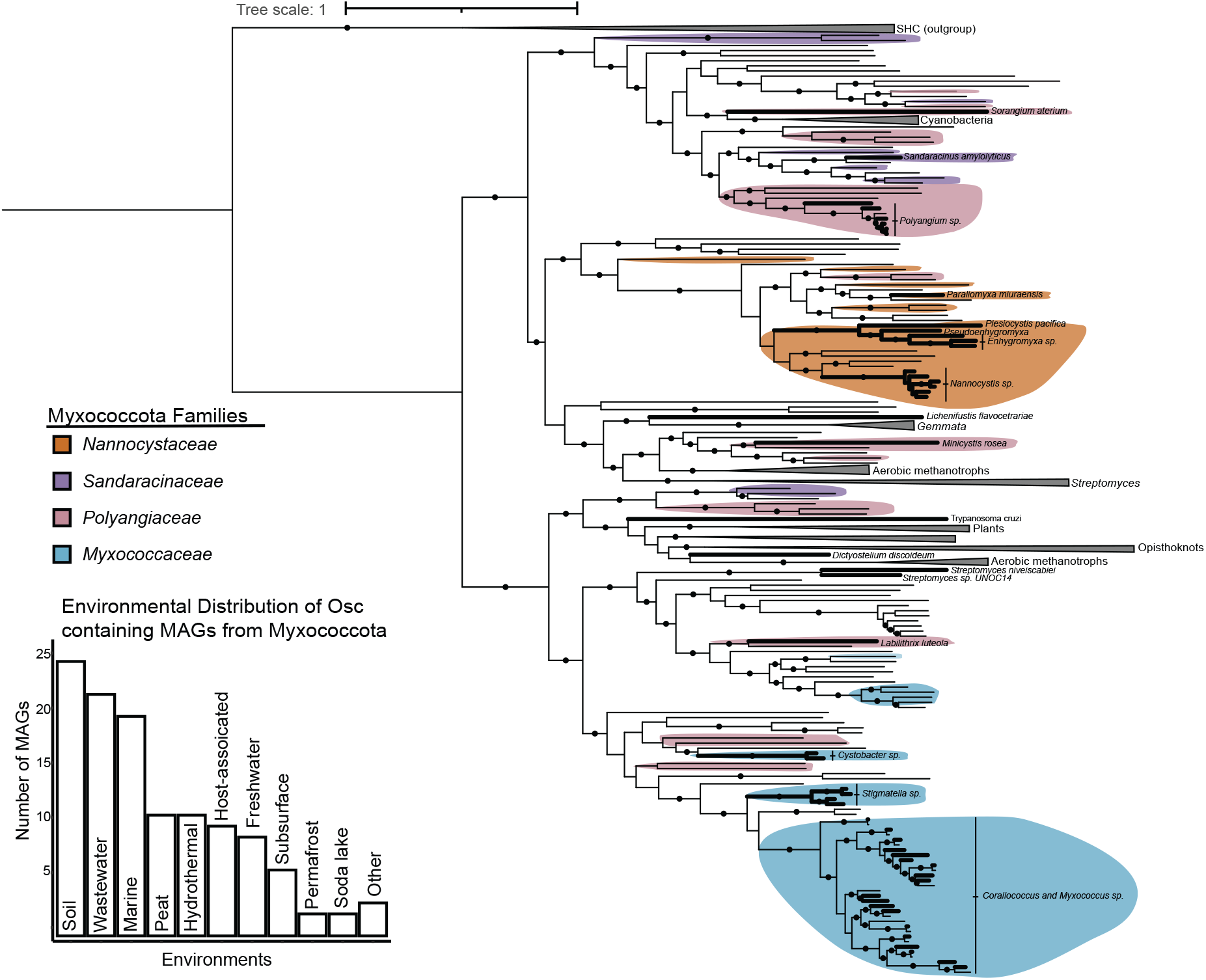
Phylogenetic analysis of uncultured myxobacterial oxidosqualene (Osc) cyclase sequences. A maximum likelihood trees of Osc sequences with squalene hopane cyclase as an outgroup was generated using IQTree with the model of best fit and 5000 ultrafast bootstrap replicates. Branches with bootstrap support values >90 are denoted with a black circle. Bolded branches represent sequences from cultured organisms. Normal weight branches represent sequences from myxobacterial metagenome assembled genomes. Sequences with a taxonomic ranking corresponding to the family level or lower are colored by family as follows: *Nannocystaceae* (orange), *Sandaracinaceae* (purple), *Polyangiaceae* (pink), and *Myxococcaceae* (blue). A bar graph representing the environmental distribution of metagenomic sequences is provided.

We performed a similar analysis for C-14 demethylase (Cyp51) sequences from Myxococcota MAGs, identifying 145 unique sequences (*SI Appendix*, Fig. S13). As with the Osc MAG sequences, many of these sequences fall into the same clades as Cyp51 sequences from cultured myxobacteria but on longer branches, exhibiting diversity missed in the current cultured bacteria. When assigned a more specific taxonomic ranking, all *Nannocystaceae* sequences except one cluster with cultured *Nannocystaceae* sequences and the uncultured *Polyangiaceae* sequences are found throughout the tree, paralleling what we observed with Osc MAG sequences. Additionally, we found Cyp51 sequences in MAGs from *Kofleriaceae* and *Vulgatibacteraceae*, two families with only a single sequenced, cultured representative that does not synthesize sterols, suggesting that we may yet find sterol synthesizers in these lineages. Taken together, these MAG sequences both illustrate unappreciated diversity in myxobacterial sterol synthesizers and further support the phylogenetic patterns we observed in sterol biosynthesis proteins from cultured myxobacteria.

## Discussion

In this study, we identified two additional myxobacteria, *S. amylolyticus* and *M. rosea*, capable of de novo cholesterol biosynthesis and find that sterol production in these bacteria differs from what we have previously observed in *E. salina* by the accumulation of only C-24 saturated sterols. This sterol profile suggests C-24 reduction occurs as in intermediary step in cholesterol biosynthesis in *S. amylolyticus* and *M. rosea* in line with the Kandutsch-Russell (K-R) cholesterol biosynthesis pathway and not the Bloch pathway, where C-24 reduction is the terminal step (34, 35). In eukaryotes, the use of these two pathways is thought to be a mechanism for regulating production of cholesterol intermediates as these lipids can serve specific physiological functions and act as precursors for downstream metabolite production (36–38). In all three of these myxobacteria, cholesterol intermediates comprise a significant portion of the sterol profile, suggesting that these intermediates may also serve specific, unrecognized physiological functions and that the differences in pathway usage may translate into distinct molecular interactions governing sterol biology in *Polyangiaceae, Sandaracinaceae*, and *Nannocystaceae*.

Underlying the differences in cholesterol intermediate production amongst these myxobacteria is our discovery of a previously unknown bacterial C-24 sterol reductase (Bsr) belonging to the geranylgeranyl reductase protein (GGR) family. The Bsr protein expands the chemistry that GGR proteins are known to perform in vivo to include substrates that are largely cyclic instead of linear, hydroxylated instead of phosphorylated, and comprised of a short isoprenoid chain. Bsr is found in all three sterol producing families of Myxococcota, however, its distribution is sporadic and, in some cases, redundant. While both *S. amylolyticus* and *M. rosea* only have the bacterial C-24 sterol reductase, *E. salina* harbors homologs to both the canonical C-24 reductase and the bacterial C-24 sterol reductase we have identified here. However, analysis of the *E. salina* proteome suggests that only the canonical C-24 reductase is translated under our current culturing conditions. We posit that the differences we have observed in the sterol profile between *E. salina* and the other cholesterol producing myxobacteria are driven by the expression of the canonical C-24 reductase over Bsr and that these two enzymes in *E. salina* may provide biosynthetic flexibility in the sterol intermediates produced. Bsr also further differentiates bacterial sterol production from eukaryotes. Bsr is only found in the genomes of sterol producing bacteria, primarily from the phyla Myxococcota and Cyanobacteriota, and not eukaryotes. This unique bacterial sterol reductase, alongside the distinct bacterial enzymes for C-4 demethylation (9, 39), highlights an evolutionary history for bacterial cholesterol biosynthesis at least partially distinct from eukaryotes, implying the pathway we see in modern myxobacteria was not acquired directly from a eukaryotic source. Indeed, complex sterol production across the bacterial domain-including species in Actinobacteria, Cyanobacteria, and aerobic methanotrophs-is characterized by sterol biosynthetic enzymes of bacterial origin, suggesting that the proliferation of this pathway in many of these bacteria was likely driven by horizontal gene transfer between bacterial species and not necessarily between eukaryotes and bacteria.

Modern day myxobacteria produce a diverse array of sterol products. Yet, the evolutionary history of sterol biosynthesis within this phylum remains unclear, obscuring if, and how, these bacteria contributed to the evolution of sterol biosynthesis in eukaryotes. As a phylum, Myxococcota is sufficiently ancient for their ancestors to have been present when the last common eukaryotic ancestor (LECA) formed, with current estimates placing the divergence of this phylum before the Great Oxidation Event (40). Thus, an ancient myxobacterial source for sterol production in LECA is plausible. However, the families that comprise Myxococcota may themselves be quite young, with molecular clock analysis of a chitinase found in members of *Myxococcaceae* placing the age of this family at 275 Mya (41), long after the divergence of crown group eukaryotes. Understanding when Myxococcota acquired the sterol biosynthesis genes we observe in modern day organisms could help constrain the role of these bacteria in the evolution of sterol biosynthesis. The phylogenetic relationship of Osc sequences and the split between cycloartenol and lanosterol synthase in this phylum indicates the capacity for sterol production was acquired at least twice, once in *Myxococcaceae* and one or more times in Polyangia. In the current cultured isolates, cycloartenol synthase is almost entirely restricted to *Myxococcaceae*, suggesting this Osc homolog was acquired after the *Myxococcaceae* diverged from *Anaeromyxobacteraceae* and *Vulgatibacteraceae*. If this is the case, it suggests cycloartenol synthase is a recent acquisition by the phylum and is unlikely to be a source for sterol biosynthesis in LECA. However, we found several cycloartenol synthase sequences from MAGs assigned to *Polyangiaceae* that cluster with the sequences from *Myxococcaceae*. This, along with the cycloartenol synthase found in *Labilithrix luetola*, may point to the potential for an earlier acquisition of cycloartenol synthase by Myxococcota.

Similarly, the genes responsible for complex sterol production display a distribution and phylogenetic pattern that makes inferring sterol ancestry in Myxococcota difficult. For example, complex sterol production appears ancestral in *Nannocystaceae* but is not found in the single cultured isolate that makes up the closely related family *Kofleriaceae*. Additionally, the proteins necessary for downstream sterol modifications appear sparsely in *Polyangiaceae*, but gene loss could be a significant factor in this family, as we likely observed in the *Polyangium* species. We also find examples of specific proteins, such as the Cyp51 sequences in MAGs, assigned to myxobacterial taxonomic groups where we do not otherwise find sterol biosynthesis genes, suggesting a capacity for complex sterol production may be more universal in Myxococcota and providing support for a more ancient acquisition of this pathway. Ultimately, further culturing and sequencing coupled to phylogenetic and molecular analyses is needed to provide the nuanced understanding of sterol biosynthesis within Myxococcota required to resolve these broader evolutionary questions.

Finally, while phylogenomic analyses of bacterial sterol biosynthesis proteins have provided insight into the evolution of these lipids, the molecular mechanisms involved in bacterial sterol function could provide further information about the evolutionary history of these molecules in bacteria. Currently, our understanding of bacterial sterol physiology is limited to studies of essentiality in two bacteria (8, 42, 43). These studies provide a foundation for exploring the function of sterols in bacteria and have highlighted specific cellular processes in which sterols may play an important role. However, we still have little insight into the specific proteins involved in bacterial sterol regulation, transport, or broader physiology. In eukaryotes, protein-sterol interactions are central to a range of different processes, including regulation of lipid biosynthesis, trafficking between organelles, and triggering signaling pathways (44–49). These interactions can be, to varying extents, conserved. For example, sterol regulatory element-binding proteins (SREBPs) are a family of transcription factors, themselves regulated by the sterol SREBP cleavage activating proteins (SCAPs) (50), responsible for modulating endogenous cholesterol production in animals (51) that have also been implicated in lipid and hypoxia regulation in yeast (52, 53). However, SREBP homologs are not present in sterol-producing myxobacterial genomes, or any bacterial genomes, indicating that alternative pathways for sterol regulation may occur in bacteria. Identifying how Myxococcota species regulate and localize sterols could provide insight into the conservation of these interactions across the phylum, providing a more accurate view of both sterol biology in these bacteria and the evolutionary relationships underlying the synthesis of these important lipids across all domains of life.

## Methods

### Bacterial Cell Culture

*Sandaracinus amylolyticus* DSM 53668 and *Minicystis rosea* DSM 24000 were cultured in a minimal media (wt/vol: 0.05% calcium chloride dihydrate, 0.01 % magnesium sulfate heptahydrate, and 25 mM HEPES buffer) adjusted to pH 7 and supplemented with autoclaved whole cell *Escherichia coli*. Cultures were grown for 10 days and fed 7ml of supplemental *E. coli*, in 20ml liquid cultures shaking at 220 rpm, at 30 °C. Supplemental concentrated *E. coli* was prepared by inoculating a 500 ml LB culture with 500 µl of an overnight DH10B *E. coli* culture and grown for 18hr at 37 °C, shaking at 220 rpm. *E. coli* cells were then harvested by centrifugation, resuspended in 50ml of the minimal myxobacteria media, and autoclaved. Lysate experiments were done using *E. coli* BL21. *E. coli* strains were cultured on Luria broth (LB) or TYGPN media (54), at 37 °C or 25 °C, shaking at 220 rpm, and supplemented with 20µg/ml of chloramphenicol and 30µg/ml of kanamycin, as necessary.

### Molecular Cloning Techniques

Plasmids and oligonucleotides used in this study are described in *SI Appendix* Table S3 and S4. Oligonucleotides were purchased from Integrated DNA Technologies (Coralville, IA). Genomic DNA from *S. amyloloticus* was isolated using the GeneJET Genomic DNA Purification Kit (Thermo Scientific). Plasmid DNA was isolated using the GeneJET Plasmid Miniprep Kit (Thermo Scientific). DNA fragments used during cloning were isolated using the GeneJET Gel Extraction Kit (Thermo Scientific). DNA was sequenced by ELIM Biopharm (Hayward, CA).

The expression plasmid used in cell free lysate experiments was constructed by sequence and ligation independent cloning (SLIC) (55). Briefly, gene fragments were amplified from genomic DNA using Phusion DNA Polymerase (New England Biolabs) with complementary plasmid insertion site overhangs and gel purified. The expression plasmids were linearized by digestion. Complementary overhangs were created on both gel-purified gene fragments and linearized plasmids by incubation with T4 DNA Polymerase (MilliporeSigma) without nucleotides. Vectors and gene fragments were then annealed and transformed by electroporation without ligation into *E. coli. E. coli* strains were transformed by electroporation using a MicroPulser Electroporator (BioRad) as recommended by the manufacturer.

### Cell Free Lysate Experiments

Cultures were grown in 20ml of TYPGN media at 37°C, shaking at 220rpm until an optical density (OD) of 0.5 was reached. Expression was induced with 100 µM isopropyl β-d-1-thiogalactopyranoside (IPTG). After induction, cultures were grown at 25°C, shaking at 220rpm for 18 hours. Cells were harvested by centrifugation at 4,500xg for 10 minutes and resuspended in lysis buffer (50mM Tris buffer, 200mM sodium chloride, and 65 mM dithiothreitol at pH 7.5), as previously described for geranylgeranyl reductase in vitro experiments (26). Cells were lysed by sonication with pulses at an amplitude of 30% for 5 seconds on, 15 seconds off, for 4 minutes. Lysates were clarified by centrifugation at 14,000xg for 15 minutes. 200µM flavin adenine dinucleotide (FAD) and 100µM of either desmosterol or zymosterol was added to lysate for a total reaction volume of 500 µl. Reactions were incubated for 18 hours at 30°C before lipid products were extracted.

### Lipid extractions

Myxobacterial cultures were spun down, lyophilized, and massed before sterol analysis. Lipids were extracted from dried cell pellets using a modified Bligh Dyer extraction (56). Briefly, cells were resuspended in 10: 5: 4 methanol: dichloromethane (DCM): water (vol: vol: vol) and sonicated in a water bath for 1 hour. Samples were then phase separated by adding 2 volumes of 1:1 water and dichloromethane and the organic layer was separated by centrifugation, removed, and dried down under N_2_. This total lipid extract was then analyzed using gas chromatography-mass spectrometry (GC-MS).

### GC-MS

Lipids were derivatized to trimethylsilyl ethers in 1:1 (vol: vol) Bis(trimethylsilyl)trifluoroacetamide: pyridine by heating to 70 °C for 1 h and analyzed on an Agilent 7890B Series GC coupled to a 5977 A Series MSD. 2 μL of each sample was injected in splitless mode at 250 °C. Lipids were separated on a 60 m Agilent DB17HT column (60 m x 0.25 mm i.d. x 0.1 μm film thickness) with helium as the carrier gas at a constant flow of 1.1 mL/min and programed as follows: 100 °C for 2 min; then 8 °C/min to 250 °C and held for 10 min; then 3 °C/min to 330 °C and held for 17 min. The ion source was set at 230 °C and operated at 70 eV in EI mode scanning from 50-850 Da in 0.5 s. Samples were analyzed using Agilent MassHunter Qualitative Analysis (version B.06.00) and sterols identified based on retention time and spectra. Sterol spectra in samples were compared to laboratory standards and spectra deposited in the American Oil Chemists’ Society (AOCS) Lipid Library (http://lipidlibrary.aocs.org/index.cfm) or the National Institute of Standards and Technology (NIST) databases.

### Bioinformatic Techniques and Phylogenetic Analysis

Sterol biosynthesis homologs in Myxococcota were identified in cultured myxobacterial species by a BLASTp search (e-value cutoff e-50) of Myxococcota sequence in the NCBI database against the cholesterol biosynthesis proteins in *E. salina* (9, 57). A more restrictive e-value cutoff was used to identify putative sterol biosynthesis proteins because many of these proteins belong to ubiquitous superfamilies and this more conservative cutoff limits overestimation of the biosynthetic capabilities of these organisms. Sterol biosynthesis genes in uncultured Myxococcota MAGs were identified by a tBLASTn search (e-value cutoff e-50). Sequences were limited to those 400 amino acids or longer for oxidosqualene cyclase and 300 amino acids for the C-14 demethylase. Redundant metagenomic sequences were removed using the Decrease Redundancy tool (http://web.expasy.org/decrease_redundancy/).

Sequences were aligned using MUSCLE in MEGA (version 11.0.13) and exported to a FASTA file. Phylogenetic trees were generated using IQ-TREE2 (version 2.2.2.6) (58). Each tree was run with 5000 ultrafast bootstrap replicates (59). Modelfinder was used to identify the best fit model for each protein and a maximum likelihood tree was generated. Trees were also generated using the three next best fitting models according to modelfinder to test tree robustness. Phylogenetic trees were then edited for publication using the iTOL website (http://itol.embl.de/).

## References

1. J. H. Wei, X. Yin, P. V. Welander, Sterol Synthesis in Diverse Bacteria. Front Microbiol 7, 990 (2016).

2. A. Pearson, M. Budin, J. J. Brocks, Phylogenetic and biochemical evidence for sterol synthesis in the bacterium Gemmata obscuriglobus. Proceedings of the National Academy of Sciences 100, 15352–15357 (2003).

3. E. Desmond, S. Gribaldo, Phylogenomics of Sterol Synthesis: Insights into the Origin, Evolution, and Diversity of a Key Eukaryotic Feature. Genome Biology and Evolution 1, 364– 381 (2009).

4. M. Dworkin, Fibrils as extracellular appendages of bacteria: Their role in contact-mediated cell-cell interactions in Myxococcus xanthus. BioEssays 21, 590–595 (1999).

5. S. Huntley, et al., Comparative Genomic Analysis of Fruiting Body Formation in Myxococcales. Molecular Biology and Evolution 28, 1083–1097 (2011).

6. J. Muñoz-Dorado, F. J. Marcos-Torres, E. García-Bravo, A. Moraleda-Muñoz, J. Pérez, Myxobacteria: Moving, Killing, Feeding, and Surviving Together. Front. Microbiol. 7 (2016).

7. D. Wall, D. Kaiser, Type IV pili and cell motility. Mol Microbiol 32, 1–10 (1999).

8. H. B. Bode, et al., Steroid biosynthesis in prokaryotes: identification of myxobacterial steroids and cloning of the first bacterial 2,3(S)-oxidosqualene cyclase from the myxobacterium Stigmatella aurantiaca. Molecular Microbiology 47, 471–481 (2003).

9. A. K. Lee, J. H. Wei, P. V. Welander, De novo cholesterol biosynthesis in bacteria. Nat Commun 14, 2904 (2023).

10. D. Gawas, R. Garcia, V. Huch, R. Müller, A Highly Conjugated Dihydroxylated C28 Steroid from a Myxobacterium. J. Nat. Prod. 74, 1281–1283 (2011).

11. Y. Khatri, et al., Substrate Hunting for the Myxobacterial CYP260A1 Revealed New 1α-Hydroxylated Products from C-19 Steroids. ChemBioChem 17, 90–101 (2016).

12. S. G. Salamanca-Pinzon, et al., Structure–function analysis for the hydroxylation of Δ4 C21-steroids by the myxobacterial CYP260B1. FEBS Letters 590, 1838–1851 (2016).

13. S. H. Akone, J. J. Hug, A. Kaur, R. Garcia, R. Müller, Structure Elucidation and Biosynthesis of Nannosterols A and B, Myxobacterial Sterols from Nannocystis sp. MNa10993. J. Nat. Prod. 86, 915–923 (2023).

14. J. Amiri Moghaddam, et al., Analysis of the Genome and Metabolome of Marine Myxobacteria Reveals High Potential for Biosynthesis of Novel Specialized Metabolites. Sci Rep 8, 16600 (2018).

15. S. Felder, et al., Salimyxins and Enhygrolides: Antibiotic, Sponge-Related Metabolites from the Obligate Marine Myxobacterium Enhygromyxa salina. ChemBioChem 14, 1363–1371 (2013).

16. J. J. Brocks, et al., Biomarker evidence for green and purple sulphur bacteria in a stratified Palaeoproterozoic sea. Nature 437, 866–870 (2005).

17. J. J. Brocks, et al., Lost world of complex life and the late rise of the eukaryotic crown. Nature 618, 767–773 (2023).

18. D. A. Gold, A. Caron, G. P. Fournier, R. E. Summons, Paleoproterozoic sterol biosynthesis and the rise of oxygen. Nature 543, 420–423 (2017).

19. R. H. Anderson, et al., Sterols lower energetic barriers of membrane bending and fission necessary for efficient clathrin-mediated endocytosis. Cell Reports 37, 110008 (2021).

20. A. Heese-Peck, et al., Multiple Functions of Sterols in Yeast Endocytosis. MBoC 13, 2664– 2680 (2002).

21. H. Pichler, H. Riezman, Where sterols are required for endocytosis. Biochimica et Biophysica Acta (BBA) - Biomembranes 1666, 51–61 (2004).

22. C. Santana-Molina, E. Rivas-Marin, A. M. Rojas, D. P. Devos, Origin and Evolution of Polycyclic Triterpene Synthesis. Molecular Biology and Evolution 37, 1925–1941 (2020).

23. Y. Hoshino, E. A. Gaucher, Evolution of bacterial steroid biosynthesis and its impact on eukaryogenesis. Proceedings of the National Academy of Sciences 118, e2101276118 (2021).

24. P. LóPez-García, D. Moreira, “The Syntrophy Hypothesis for the Origin of Eukaryotes” in Symbiosis: Mechanisms and Model Systems, J. Seckbach, Ed. (Springer Netherlands, 2002), pp. 131–146.

25. P. López-García, D. Moreira, The Syntrophy hypothesis for the origin of eukaryotes revisited. Nat Microbiol 5, 655–667 (2020).

26. C. W. Meadows, et al., Discovery of novel geranylgeranyl reductases and characterization of their substrate promiscuity. Biotechnol Biofuels 11, 340 (2018).

27. D. Sasaki, et al., Structure and Mutation Analysis of Archaeal Geranylgeranyl Reductase. Journal of Molecular Biology 409, 543–557 (2011).

28. P. Wang, et al., Identification of a Geranylgeranyl reductase gene for chlorophyll synthesis in rice. SpringerPlus 3, 201 (2014).

29. A. Upadhyay, et al., Mycobacterial MenJ: An Oxidoreductase Involved in Menaquinone Biosynthesis. ACS Chem. Biol. 13, 2498–2507 (2018).

30. M. M. Meyer, M. J. R. Segura, W. K. Wilson, S. P. T. Matsuda, Oxidosqualene Cyclase Residues that Promote Formation of Cycloartenol, Lanosterol, and Parkeol. Angewandte Chemie International Edition 39, 4090–4092 (2000).

31. W. Kohl, A. Gloe, H. Reichenbach, Steroids from the Myxobacterium Nannocystis exedens. Microbiology 129, 1629–1635 (1983).

32. Y. Liu, Q. Yao, H. Zhu, Meta-16S rRNA Gene Phylogenetic Reconstruction Reveals the Astonishing Diversity of Cosmopolitan Myxobacteria. Microorganisms 7, 551 (2019).

33. J. Wang, J. Wang, S. Wu, Z. Zhang, Y. Li, Global Geographic Diversity and Distribution of the Myxobacteria. Microbiol Spectr 9, e00012–21 (2021).

34. K. Bloch, The Biological Synthesis of Cholesterol. Science 150, 19–28 (1965).

35. A. A. Kandutsch, A. E. Russell, Preputial Gland Tumor Sterols: III. A METABOLIC PATHWAY FROM LANOSTEROL TO CHOLESTEROL. Journal of Biological Chemistry 235, 2256–2261 (1960).

36. M. A. Mitsche, J. G. McDonald, H. H. Hobbs, J. C. Cohen, Flux analysis of cholesterol biosynthesis in vivo reveals multiple tissue and cell-type specific pathways. eLife 4, e07999 (2015).

37. A. V. Prabhu, W. Luu, D. Li, L. J. Sharpe, A. J. Brown, DHCR7: A vital enzyme switch between cholesterol and vitamin D production. Progress in Lipid Research 64, 138–151 (2016).

38. C. Yang, et al., Sterol Intermediates from Cholesterol Biosynthetic Pathway as Liver X Receptor Ligands *. Journal of Biological Chemistry 281, 27816–27826 (2006).

39. A. K. Lee, et al., C-4 sterol demethylation enzymes distinguish bacterial and eukaryotic sterol synthesis. Proceedings of the National Academy of Sciences 115, 5884–5889 (2018).

40. L. Li, et al., Globally distributed Myxococcota with photosynthesis gene clusters illuminate the origin and evolution of a potentially chimeric lifestyle. Nat Commun 14, 6450 (2023).

41. D. S. Gruen, J. M. Wolfe, G. P. Fournier, Paleozoic diversification of terrestrial chitin-degrading bacterial lineages. BMC Evol Biol 19, 34 (2019).

42. L. R. Gudde, M. Hulce, A. H. Largen, J. D. Franke, Sterol synthesis is essential for viability in the planctomycete bacterium Gemmata obscuriglobus. FEMS Microbiology Letters 366, fnz019 (2019).

43. E. Rivas-Marin, et al., Essentiality of sterol synthesis genes in the planctomycete bacterium Gemmata obscuriglobus. Nat Commun 10, 2916 (2019).

44. I. Deshpande, et al., Smoothened stimulation by membrane sterols drives Hedgehog pathway activity. Nature 571, 284–288 (2019).

45. F. A. Horenkamp, D. P. Valverde, J. Nunnari, K. M. Reinisch, Molecular basis for sterol transport by StART-like lipid transfer domains. The EMBO Journal 37, e98002 (2018).

46. J. D. Horton, Sterol regulatory element-binding proteins: transcriptional activators of lipid synthesis. Biochemical Society Transactions 30, 1091–1095 (2002).

47. C. Qi, G. Di Minin, I. Vercellino, A. Wutz, V. M. Korkhov, Structural basis of sterol recognition by human hedgehog receptor PTCH1. Science Advances 5, eaaw6490 (2019).

48. R. Sato, et al., Sterol-dependent Transcriptional Regulation of Sterol Regulatory Element-binding Protein-2 *. Journal of Biological Chemistry 271, 26461–26464 (1996).

49. Z. Xu, W. Farver, S. Kodukula, J. Storch, Regulation of Sterol Transport between Membranes and NPC2. Biochemistry 47, 11134–11143 (2008).

50. S. H. Lee, J.-H. Lee, S.-S. Im, The cellular function of SCAP in metabolic signaling. Exp Mol Med 52, 724–729 (2020).

51. D. Eberlé, B. Hegarty, P. Bossard, P. Ferré, F. Foufelle, SREBP transcription factors: master regulators of lipid homeostasis. Biochimie 86, 839–848 (2004).

52. A. L. Hughes, B. L. Todd, P. J. Espenshade, SREBP pathway responds to sterols and functions as an oxygen sensor in fission yeast. Cell 120, 831–842 (2005).

53. S. D. Willger, et al., A sterol-regulatory element binding protein is required for cell polarity, hypoxia adaptation, azole drug resistance, and virulence in Aspergillus fumigatus. PLoS Pathog 4, e1000200 (2008).

54. K. Elbing, R. Brent, Recipes and tools for culture of Escherichia coli. Curr Protoc Mol Biol 125, e83 (2019).

55. M. Z. Li, S. J. Elledge, “SLIC: A Method for Sequence- and Ligation-Independent Cloning” in Gene Synthesis, (Humana Press, 2012), pp. 51–59.

56. E. G. Bligh, W. J. Dyer, A RAPID METHOD OF TOTAL LIPID EXTRACTION AND PURIFICATION. Canadian Journal of Biochemistry and Physiology (1959). 10.1139/o59-099.

57. S. F. Altschul, et al., Gapped BLAST and PSI-BLAST: a new generation of protein database search programs. Nucleic Acids Research 25, 3389–3402 (1997).

58. B. Q. Minh, et al., IQ-TREE 2: New Models and Efficient Methods for Phylogenetic Inference in the Genomic Era. Molecular Biology and Evolution 37, 1530–1534 (2020).

59. B. Q. Minh, M. A. T. Nguyen, A. von Haeseler, Ultrafast Approximation for Phylogenetic Bootstrap. Molecular Biology and Evolution 30, 1188–1195 (2013).

