## Supplemental Information for "A geranylgeranyl reductase homolog required for cholesterol production in Myxococcota"

**This PDF file includes:**

Figures S1 to S13
Tables S1 to S4
SI References

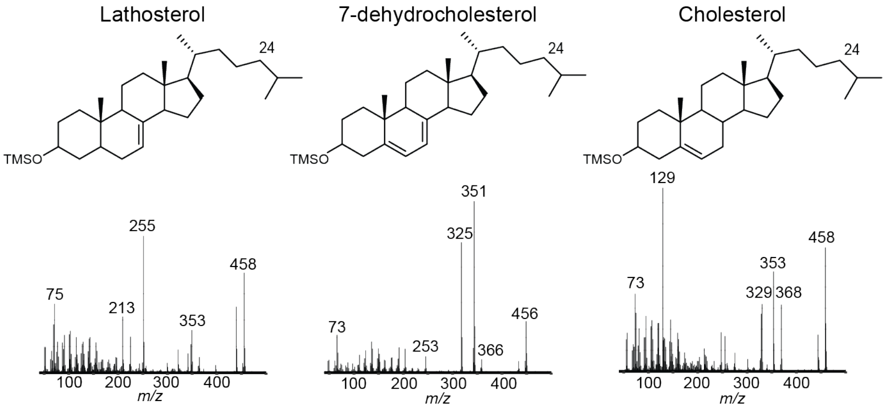


**Figure S1. Spectra of sterols identified in *Sandaracinus amylolyticus* and *Minicystis rosea* lipid extract.** Sterols were derivatized to trimethylsilyl (TMS) ether groups and separated using an Agilent 7890B series gas chromatograph with a 60m Agilent DB17 column (60m x 0.25 mm i.d x 0.1 µm film thickness). Helium gas was used as a carrier and coupled to a 5977A series mass spectrometer. Sterols were identified based on elution time and spectra compared to published spectra in the NIST database. See Methods for full extraction and GC-MS details.

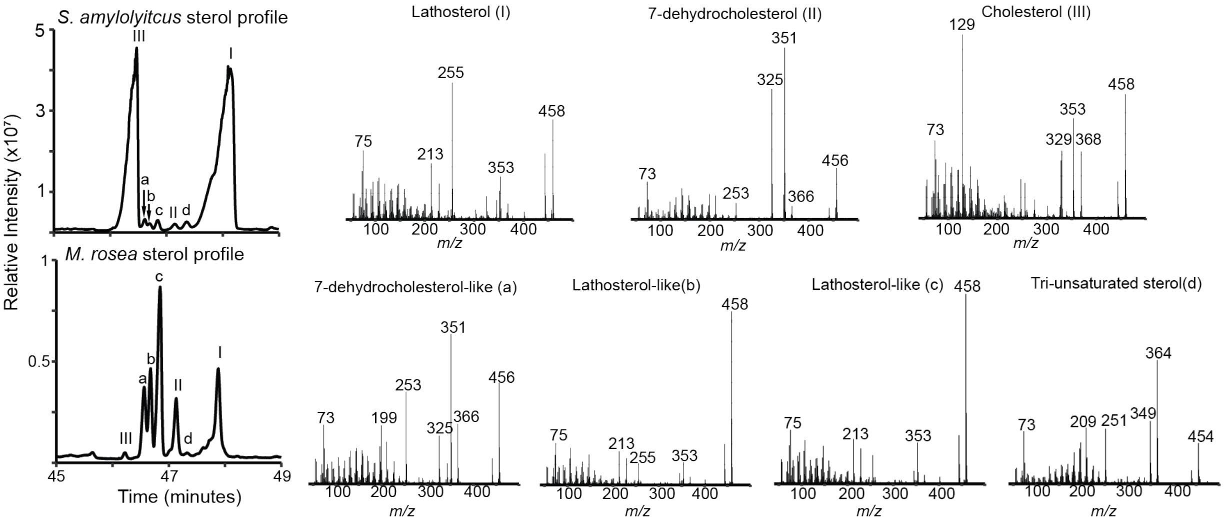


**Figure S2. Unidentified saturated sterol in *Minicystis rosea* and *Sandaracinus amylolyticus*.** Total ion chromatograms (TIC) of the lipids extracted from *M. rosea* and *S. amylolyticus*. Sterols were derivatized to trimethylsilyl ethers and identified based on spectra and elution time compared to known standards and published spectra in the NIST database. Unidentified sterols (a-d) are spectrally similar to lathosterol and 7-dehydrocholesterol but elute differently.

**Table S1. Sterol concentrations in *Sandaracinus amylolyticus* and *Minicystis rosea*.** Cell pellets from three biological replicates were lyophilized and massed before extraction. Absolute sterol concentrations were calculated based on a standard curve of cholesterol and lathosterol (20-100 ng) and normalized to pregnenolone (40ng) which was used as an internal standard. Saturated sterol a-d refers to sterols in Figure S2. BQL stands for below quantification limit.

| | ***S. amylolyticus*** | | | ***M. rosea*** | | |
|---|---|---|---|---|---|---|
| Biomass extracted (mg) | 28 | 24 | 26 | 9 | 12 | 15 |
| Total lipids extraction (mg) | 3.07 | 2.25 | 3.89 | 0.46 | 1.42 | 2.27 |
| Absolute sterol concentrations (ng/mg dry weight) | | | | | | |
| Cholesterol | 93.6 | 132.8 | 389.1 | 84.5 | 56.4 | 47.6 |
| Lathosterol | 147.5 | 338.0 | 799.2 | 315 | 107 | 76.0 |
| Relative sterol concentrations (normalized peak area/mg dry weight) | | | | | | |
| Saturated sterol a | BQL | 0.02 | 0.03 | 0.56 | 0.06 | 0.06 |
| Saturated sterol b | BQL | BQL | BQL | 0.307 | .095 | 0.065 |
| Saturated sterol c | BQL | BQL | BQL | 0.46 | 0.17 | 0.15 |
| 7-dehdyrocholesterol | BQL | BQL | BQL | 0.263 | 0.0712 | 0.058 |
| Saturated sterol d | BQL | BQL | BQL | 0.028 | 0.008 | 0.007 |

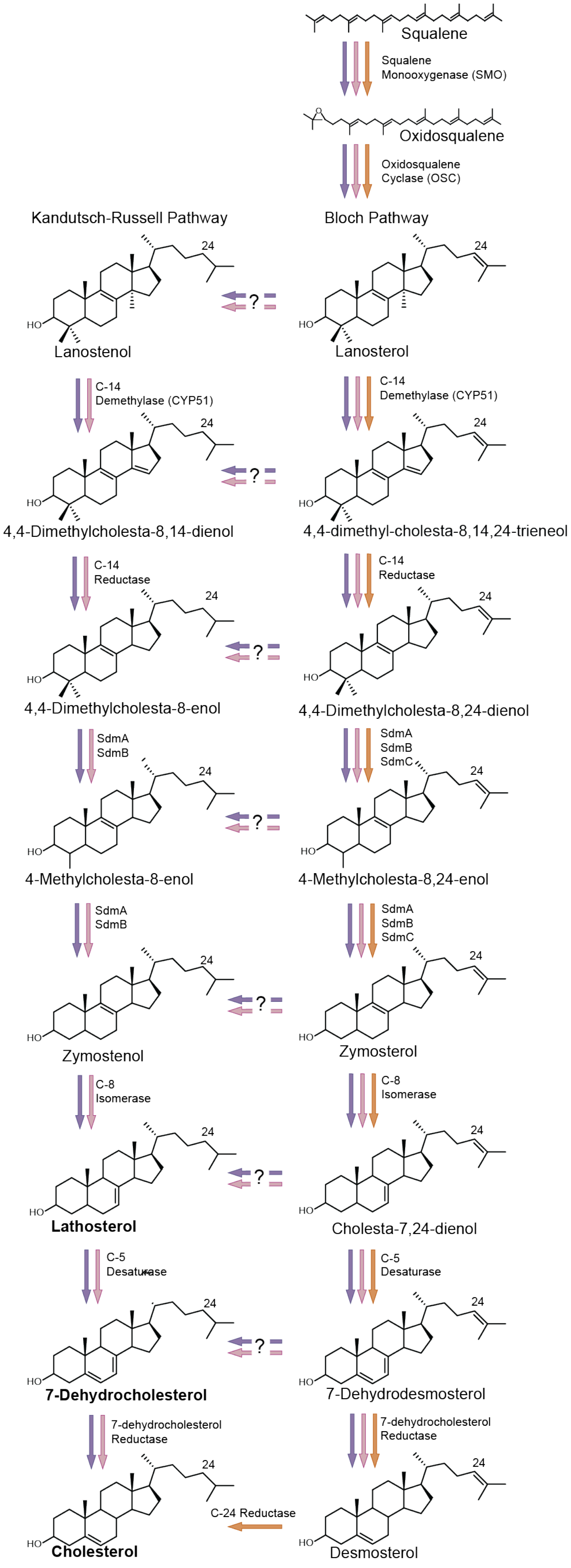


**Figure S3. The cholesterol biosynthesis pathway in *M. rosea* and *S. amylolyticus*.** A BLASTp search (e-value < $e^{-50}$) against the cholesterol biosynthesis proteins from *E. salina* was used to identify putative cholesterol biosynthesis proteins in *M. rosea* and *S. amylolyticus*. Steps present in *M. rosea* and *S. amylolyticus* are labeled with protein names and represented by pink and purple arrows, respectively. Steps with no known homolog to either the proteins present in *E. salina* or eukaryotes are labeled with a question mark. For comparison, the pathway present in *E. salina* is represented by orange arrows. Sterols detected in *M. rosea* and *S. amylolyticus* are bolded.

**Table S2. Bacteria that harbor a bacterial C-24 reductase (bcr) homolog.** The *S. amyloloticus* Bcr homolog was used in a BLASTp search to identify homologs in other genomes. The IMG locus tag of NCBI accession code for each homolog is provided as well as the e-value and percent identity.

| Organism | Locus Tag | E-value | Percent Identity |
|---|---|---|---|
| *Sandaracinus amylolyticus* | Ga0325232_116633 | 0.0 | 100% |
| *Minicystis rosea* | Ga0175540_117938 | 2e-102 | 46% |
| *Polyangium aurulentum* | Ga0588680_01_7003856_7004974 | 7e-163 | 62% |
| *Labilithrix luteola* | Ga0098217_117389 | 2e-103 | 44% |
| *Enhygromyxa salina* SWB005 | Ga0334633_116_5868_6977 | 1e-135 | 52% |
| *Enhygromyxa salina* SWB007 | Ga0334634_004_42442_43587 | 2e-147 | 55% |
| *Enhygromyxa salina* DSM 15201 | Ga0097779_110747 | 5e-142 | 54% |
| *Pseudenhygromyxa* sp. WMMC2535 | Ga0440217_01_9477840_9478970 | 8e-142 | 54% |
| *Pseudenhygromyxa* sp. WMMC2535 | Ga0440217_01_6617_7474 | 1e-97 | 52% |
| *Cystobacter fucus* | D187_008000 | 5e-137 | 56% |
| *Cystobacter ferrugineus* | Ga0248592_10182 | 1e-136 | 56% |
| *Lichenifustis flavocetrariae* | M8523_17280 | 1e-125 | 51% |
| *Nonomuraea monospora* | GCM10009850_075510 | 3e-66 | 41% |
| *Nonomuraea* sp. PA05 | Ga0444075_093_42355_43518 | 2e-80 | 41% |
| *Nostoc* sp. NIES-4103 | Ga0263809_12497 | 7e-60 | 34% |
| *Calothrix* sp. NIES 4105 | Ga0263810_115384 | 9e-17 | 24% |
| *Calothrix* sp. NIES-4107 | Ga0263900_115385 | 9e-17 | 24% |
| *Calothrix rhizoleniae* | Ga0265390_116814 | 1e-57 | 34% |
| *Amazonocrinis niriterrae* | Ga0489618_102_72015_73109 | 2e-59 | 34% |
| *Nannocystis exedens* | Ga0248598_112052 | 3e-51 | 32% |
| *Nannocystis pusilla* | Ga0576828_15_138390_139541 | 5e-48 | 33% |
| *Nannocystis radixulma* | Ga0629666_08_475685_476836 | 3e-48 | 33% |
| *Nannocystis poenicansa* | Ga0628302_01_1880090_1881145 | 7e-43 | 32% |
| *Paraliomyxa miuraensis* | Ga0617347_131_48156_49301 | 2e-47 | 36% |

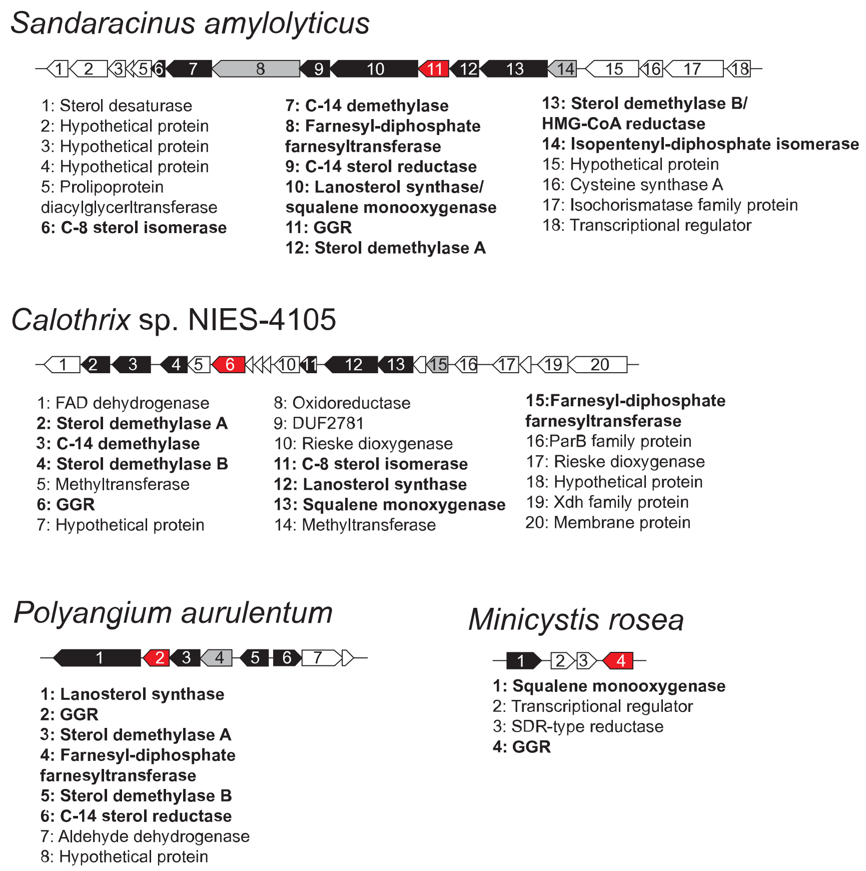


**Figure S4. Geranylgeranyl reductase co-localizes with known sterol biosynthesis genes in several bacterial genomes.** Gene clusters from selected bacteria that contain known sterol biosynthesis genes (black), upstream isoprenoid genes (grey), and the geranylgeranyl reductase homolog responsible for bacterial sterol C-24 reduction.

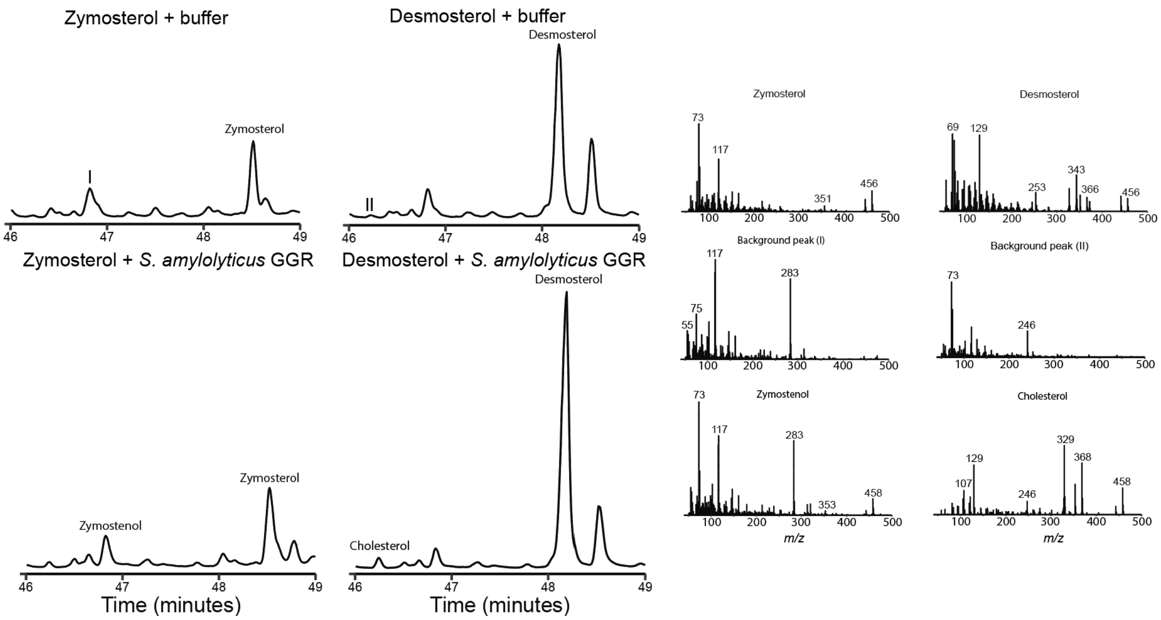


**Figure S5. Total ion chromatograms of sterol products from Bsr lysate experiments.** The substrates zymosterol and desmosterol were incubated with buffer alone (50mM Tris, 200mM NaCl, 5% glycerol, 65 mM DTT) or with lysate from *E. coli* strains overexpressing the Bsr homolog from *S. amylolyticus*. The spectra for the substrates, products and coeluting peaks are provided.

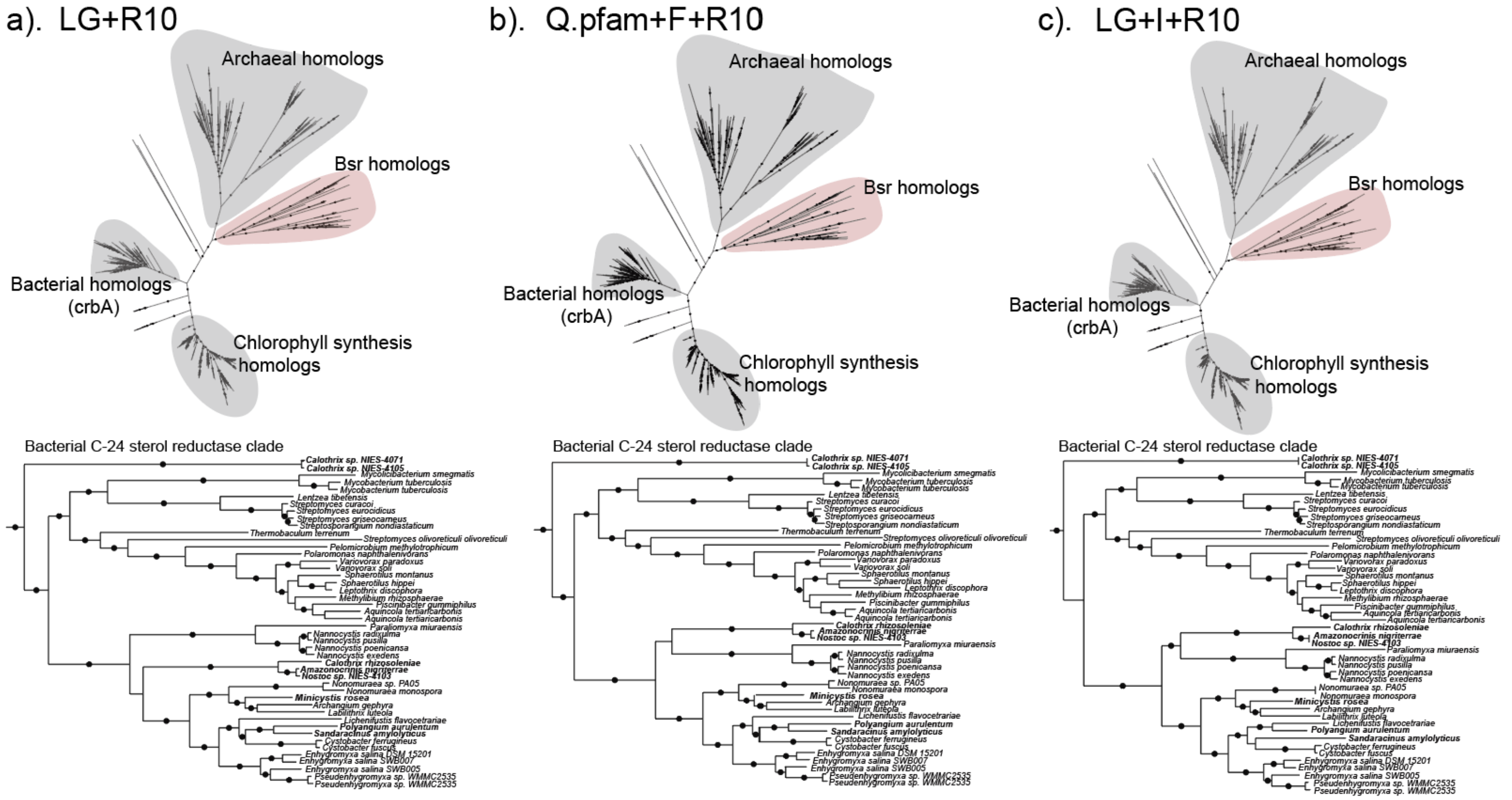


**Figure S6. Phylogenetic trees of geranylgeranyl reductase (ggr) generated using alternative models.** IQ-TREE was used to generate maximum likelihood trees with 5000 ultrafast bootstrap replicates and the following top-ranked models: a) LG+R10, with a Bayesian information criterion (BIC) score of 419746.106. b) Q.pfam+F+R10, with a BIC score of 419750.663 c) LG+I+R10, with BIC score 419791.719. Both general tree topology and the topology of the clade containing bacterial C-24 sterol reductase sequences is maintained regardless of the model used.

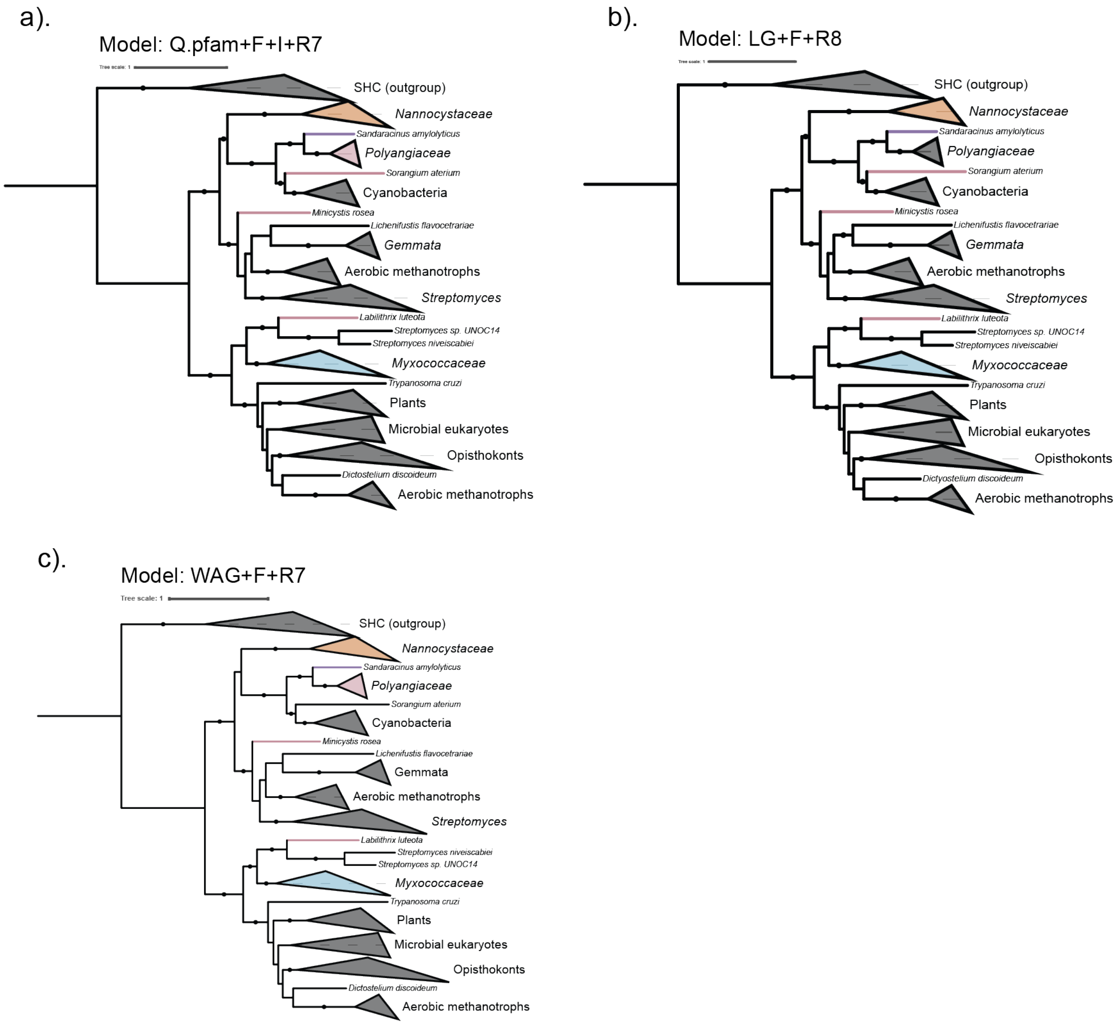


**Supplementary Figure 7. Phylogenetic trees of oxidosqualene cyclase generated using alternative models.** IQ-TREE was used to generate maximum likelihood trees with 5000 ultrafast bootstrap replicates and the following top-ranked models: a) Q.pfam.F+I+R7, with a Bayesian information criterion (BIC) score of 229166.631. b) LG+F+R8, with a BIC score of 229374.435 c) WAG+F+R7, with BIC score 229968.000. General tree topology is maintained regardless of the model used. Clades belonging to *Nannocystaceae* are colored orange, clades belonging to *Polyangiaceae* are colored pink, clades belonging to *Sandaracinaceae* are colored purple, and clades belonging to *Myxococcaceae* are colored blue.

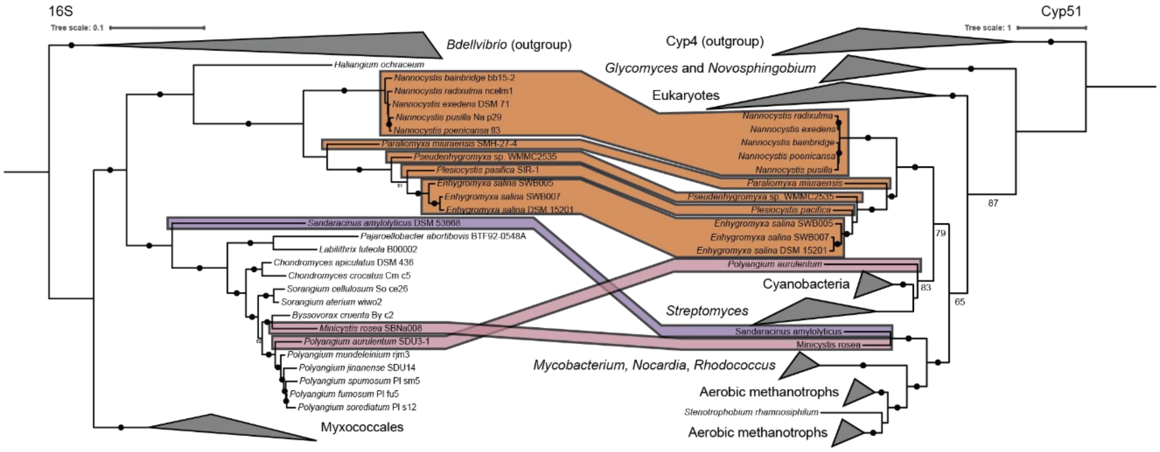


**Supplementary Figure 8 Phylogenetic analysis of myxobacterial C-14 demethylase (Cyp51) sequences in relation to taxonomic diversity.** A maximum likelihood trees of 16S sequences from Myxococcota species with *Bdellvibrio* sequences as an outgroup and Cyp51 sequences with Cyp4 sequences as an outgroup were generated using IQTree with the model of best fit and 5000 ultrafast bootstrap replicates. Branches with bootstrap support values >90 are denoted with a black circle. Myxobacterial genera are mapped from the 16S to Cyp51 tree by boxes and these boxes are colored by their respective families as follows: *Nannocystaceae* (orange), *Sandaracinaceae* (purple), and *Polyangiaceae* (pink).

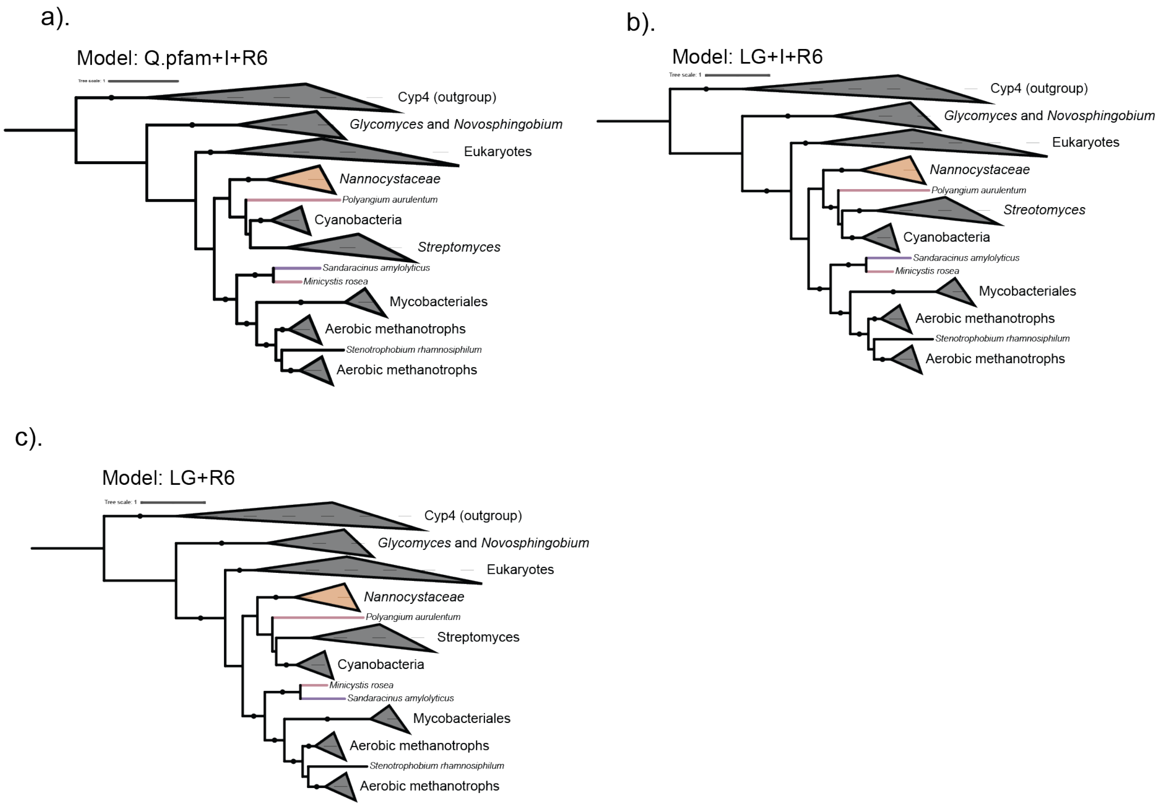


**Supplementary Figure 9. Phylogenetic trees of the C-14 demethylase (CYP51) generated using alternative models.** IQ-TREE was used to generate maximum likelihood trees with 5000 ultrafast bootstrap replicates and the following models: a) Q.pfam+I+R6 with a Bayesian information criterion (BIC) score of 109548.101. b) LG+I+R6 with BIC score of 109642.973. c) LG+R6 with a BIC score of 109643.107. General tree topology is maintained regardless of the model used. Clades belonging to *Nannocystaceae* are colored orange, clades belonging to *Polyangiaceae* are colored pink, and clades belonging to *Sandaracinaceae* are colored purple.

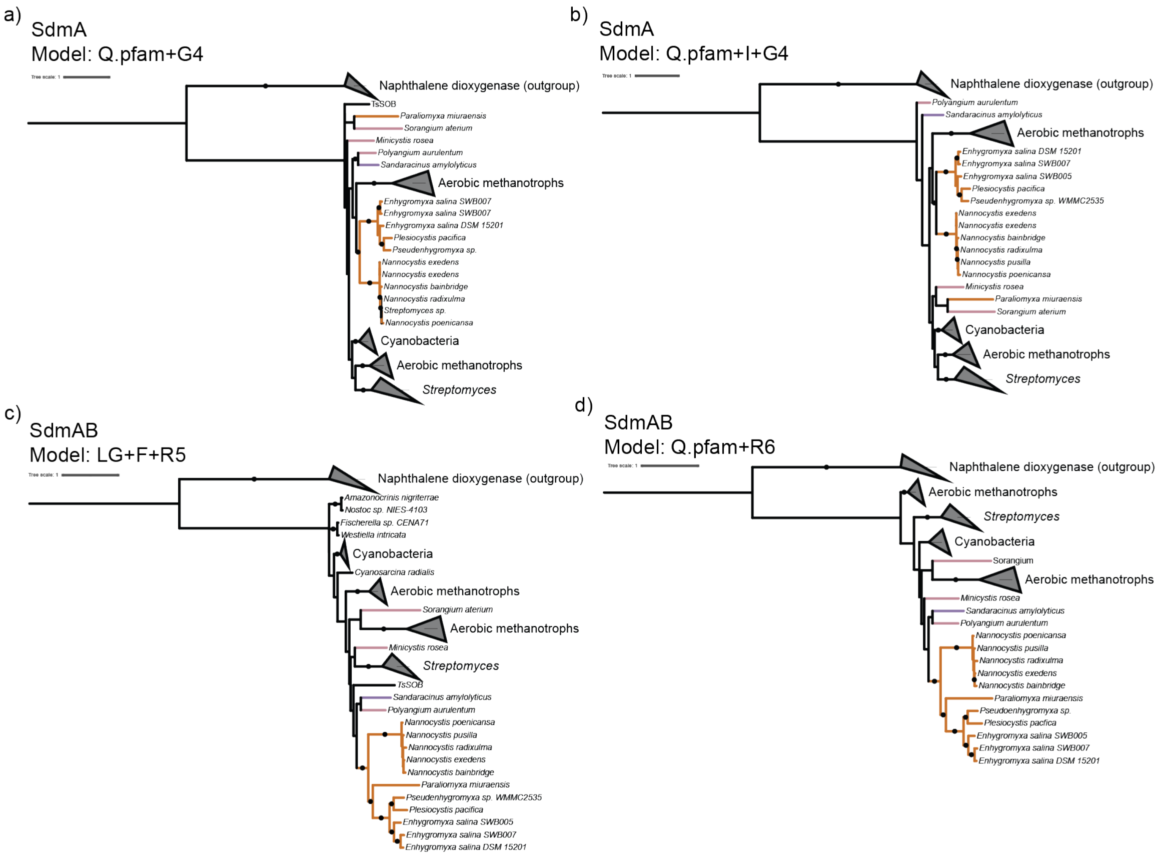


**Figure S10. Phylogenetic trees of the C-4 sterol demethylase A and B (SdmAB) generated with different models.** IQ-TREE was used to generate maximum likelihood trees with 5000 ultrafast bootstrap replicates and the following models: a) aligned SdmA sequences modeled with Q.pfam+G4, with a Bayesian information criterion (BIC) of 43063.697. b) aligned SdmA sequences modeled with Q.pfam+I+G4, with a BIC of 43064.961. c) aligned, concatenated SdmAB sequences modeled with LG+F+R5 with a BIC of 64700.937. d) aligned, concatenated SdmAB sequences modeled with Q.pfam+R6 with a BIC of 64812.153. Branches belonging to *Nannocystaceae* are colored orange, *Polyangiaceae* pink, and *Sandaracinaceae* purple.

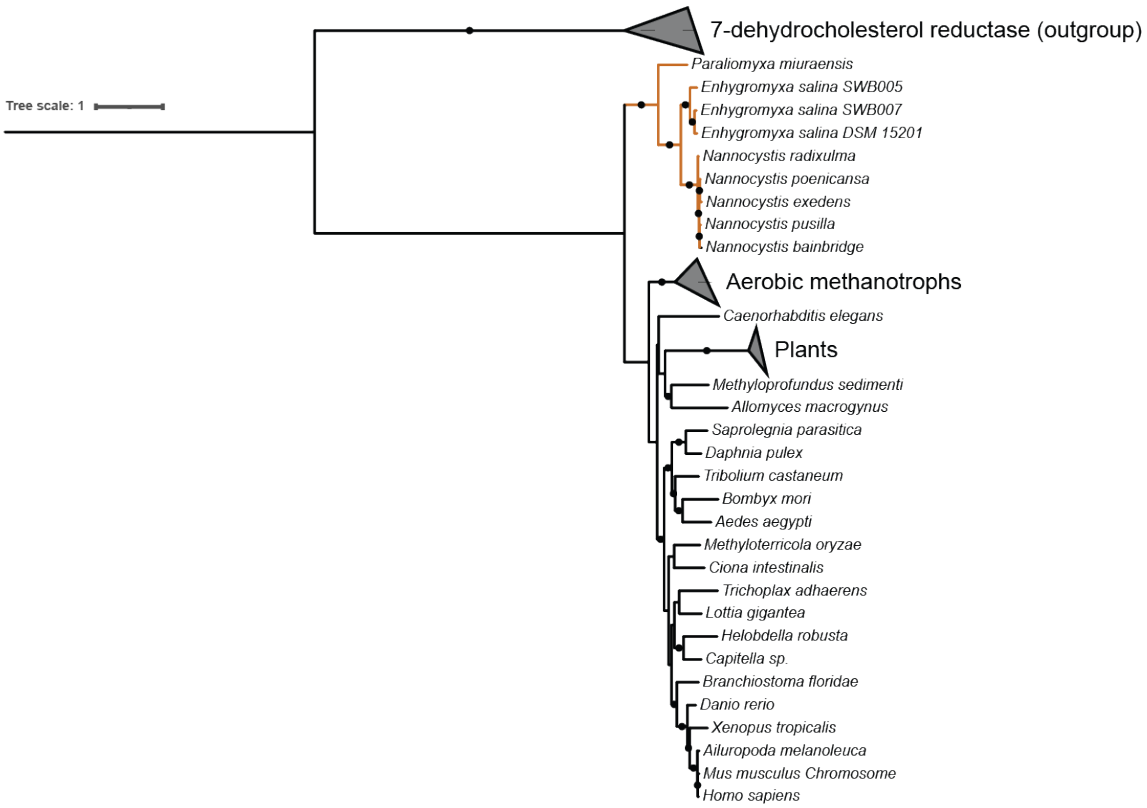


**Figure S11. Phylogenetic analysis of C-24 reductase (DHCR24) protein sequences.** IQ-TREE was used to generate a maximum likelihood trees from DHCR24 sequences with 5000 ultrafast bootstrap replicates and using model Q.pfam+G4. Branches with bootstrap support values >90 are denoted with a black circle. Branches corresponding to sequences from *Nannocystaceae* are colored orange.

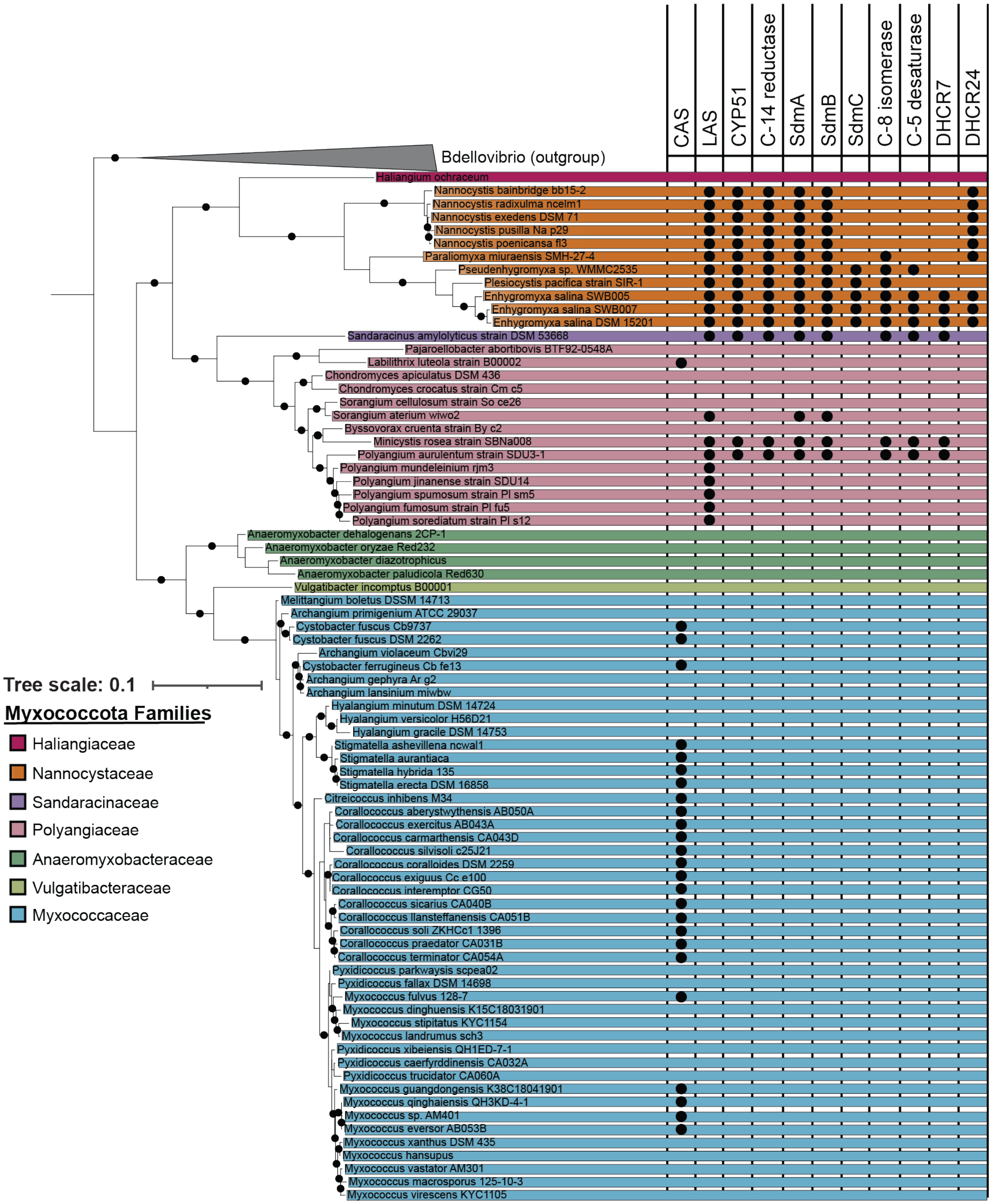


**Figure S12. Distribution of sterol biosynthesis proteins in Myxococcota.** 16S sequences of cultured myxobacteria with sequenced genomes were aligned using MUSCLE. This alignment was used to generate a maximum likelihood tree using IQ-TREE with 5000 ultrafast bootstrap replicates and the TIM2+F+I+R4 model. Sterol biosynthesis proteins were identified by a BLASTp search (e-value cutoff e-50) against the *E. salina* cholesterol biosynthesis proteins.

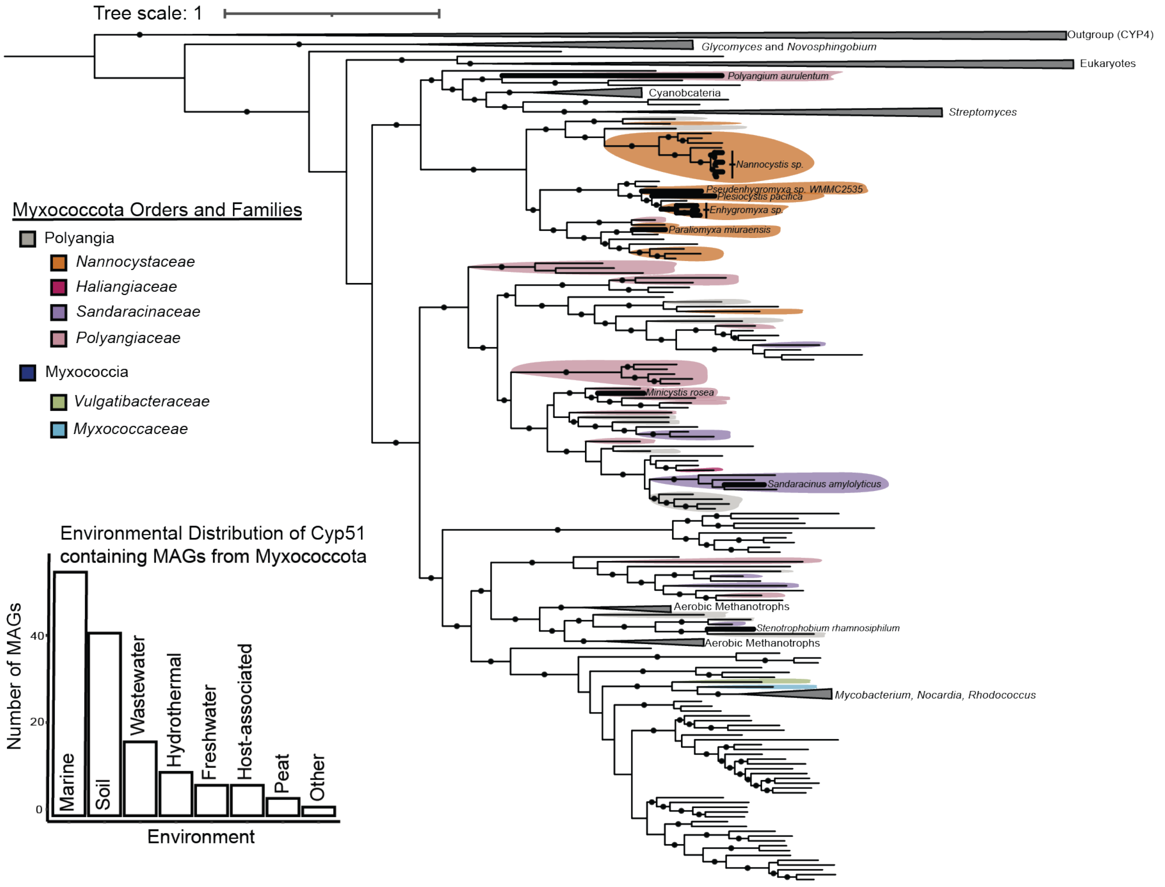


**Figure S13 Phylogenetic analysis of uncultured myxobacterial C-14 demethylase (Cyp51) sequences.** A Maximum likelihood trees of Cyp51 sequences with Cyp4 as an outgroup was generated using IQTree with the model of best fit and 5000 ultrafast bootstrap replicates. Branches with bootstrap support values >90 are denoted with a black circle. Bolded branches represent sequences from cultured organisms. Normal weight branches represent sequences from myxobacterial metagenome assembled genomes. Sequences with a taxonomic ranking corresponding to the order level or lower are colored as follows Polyangia (tan), *Nannocystaceae* (orange), *Sandaracinaceae* (purple), *Polyangiaceae* (pink), Myxococcia (indigo), *Vulgatibacteraceae* (green) *Myxococcaceae* (blue). A bar graph representing the environmental distribution of metagenomic sequences is provided.

**Table S3. Oligonucleotides used in this study.** Fwd indicates forward primer. Rv indicates reverse primer. Ga0325232_116633 indicates *Sandaracinus amylolyticus* bacterial C-24 reductase homolog.

| Oligonucleotide | Sequence | Notes |
|---|---|---|
| AL420 | ggaaaacctgtacttccaatccaat ATGGCAGAGAAGATCGACGCGAACG | SLIC pET-His10-Ga0325232_116633 Fwd |
| AL421 | ttcggatccgttatccacttccaat TCACATCTGCGCCATCAGGCG | SLIC pET-His10-Ga0325232_116633 Rv |
| AL12 | ATGCGTCCGGCGTAGAGGATC | pET28a-His 10 seq fwd |
| AL13 | GCTAGTTATTGCTTCAGAGGTGG | pET28a-His 10 seq rv |

**Table S4. Plamsids used in this study.** SAM indicates *Sandaracinus amylolyticus*. Bcr indicates the bacterial C-24 reductase. (*) indicates plasmids generated in this study.

| Plasmid | Description | Reference |
|---|---|---|
| pET-28a-His10 | T7 expression vector, modified pET28 with 10Xhis N-terminal tag | (1) |
| pACYC-GroEL/ES-TF | Co-expression copies of GroEL/ES and Trigger Factor | (2) |
| pET-28a-His10-SAM-Bcr | Ga0325232_116633 expression plasmid.<br><br>Ga0325232_116633 was amplified by PCR with primers AL420 and AL421. The fragment was assembled by SLIC into the NdeI site of pET28a-His10. Sequence was confirmed with oligos AL12 and AL13. | (*) |

**References:**


1. A. B. Banta, *et al.*, Structure of the RNA Polymerase Assembly Factor Crl and Identification of Its Interaction Surface with Sigma S. *Journal of Bacteriology* **196**, 3279–3288 (2014).

2. J. W. Lamppa, S. A. Tanyos, K. E. Griswold, Engineering *Escherichia coli* for soluble expression and single step purification of active human lysozyme. *Journal of Biotechnology* **164**, 1–8 (2013).
